# Multi-modal *in situ* profiling of human cells using the AVITI24 platform characterizes single-cell responses to acute stress

**DOI:** 10.64898/2026.10.28.762240

**Authors:** Olivia W Lang, Anne M. Gardner, Catherine M Smugereski, Amy M Lyndaker, B Franklin Pugh, William KM Lai

## Abstract

Cells begin to respond to stress within minutes. But how this early response depends on cell type, neighboring cells, or cell cycle phase has been difficult to determine, because bulk assays average across cells and synchronization can induce stress. We measured 338 transcripts, 48 proteins and phosphoproteins, and six cell paint stains in situ in 1.18 million single cells from four human cell lines exposed to H2O2, alone and in co-culture, and in A549 cells exposed to TNF-α, cadmium, dithiothreitol or lipopolysaccharide. In A549 cells, five distinct stressors changed the same targets in the same direction within 15 min, differing mainly in magnitude, and modestly increased JNK phosphorylation. By 30 min of H2O2 treatment, immediate-early genes (FOS, JUN, DDIT3, DUSP1, NR4A1) were induced most strongly in HCT116 and A549 cells. These changes occurred across the whole cell population rather than in a responsive subset. Cell line-specific differences persisted in co-culture, and the response of MCF7 cells did not increase with the number of adjacent, strongly-responding A549 cells. Ordering cells along the cell cycle from the same measurements identified 106 cycle-associated transcripts, and the response varied more between cell lines than among cell cycle phases. Overall, the measured regulators respond to stress within 30 minutes through a largely shared program whose magnitude depends more on cell identity than on cell cycle state or neighboring cell type.

## Introduction

The reliance of biochemical assays on bulk cell populations has limited our understanding of regulatory processes in individual cells, which are often partly stochastic [1]. Widely used assays such as RNA-seq (expression profiling), ATAC-seq (chromatin state) and ChIP-seq/exo (protein binding) average signal over many cells and can obscure differences among individual cells [1–4]. Understanding how individual cells maintain homeostasis under changing conditions requires assays that measure single cells.

Single-cell genomic approaches have shown that even seemingly homogeneous samples, such as cell lines, contain cells with distinct gene expression profiles that bulk methods average out [2, 5]. Because the behavior of a cell can also depend on its surroundings, spatial methods such as MERFISH, CosMx, Xenium and GeoMx additionally record the position of each cell or tissue region, and some also measure proteins [6–9]. However, few approaches measure RNA, protein and cell morphology in the same cell. Without all three, it is difficult to relate the full response of a cell to its state, such as its position in the cell cycle, and to its surroundings.

The acute stress response shows why such measurements are needed. Cells begin to respond to stress within minutes, the response varies at several levels [10], and cell types differ in how they respond, so separating shared responses from cell-type-specific ones is necessary. Within a population of one cell type, pre-existing differences between cells, such as where a cell is in the cell cycle, can determine how each cell responds [11, 12]. Within a population of one cell type, pre-existing differences between cells can determine how each cell responds [13, 14]. For example, the response could depend on where a cell is in the cell cycle. However, bulk studies of the cell cycle require synchronizing the population by double-thymidine block, nocodazole arrest, or CDK inhibition, and these treatments can themselves induce stress responses [15–17].

Several basic questions about the acute stress response remain open. It is unclear to what extent the earliest responses to chemically distinct stresses are shared or stressor-specific, and how their magnitude differs among cell types [18–20]. It is also unknown whether responses occur uniformly across a population or only in a responsive subset of cells. Additionally, a response may be intrinsic to a cell’s current state, and it may also depend on the cell’s environment. For example, responding cells could transmit a stress signal to their neighbors through paracrine or contact-dependent signals, as proposed for the bystander effects of oxidative and genotoxic stress [21–23]. Whether the response depends on cell-cycle state has also been difficult to test without synchronization [24–26].

The AVITI24 platform (Element Biosciences) allows these questions to be tested directly because it quantifies RNA, protein, and cell morphology *in situ* in the same cell [27]. Cell morphology is measured with a multiplexed cell paint stain of the cell membrane, nucleus, actin cytoskeleton, endoplasmic reticulum, Golgi apparatus, and mitochondria. RNA molecules are detected with target-specific probes and proteins with DNA-barcoded antibodies. Each measurement is quantified in two compartments, the nucleus and the whole cell. Because the spatial position of every cell is retained, the response of each cell can be compared with that of its neighbors. The probe panel used here also includes many transcripts whose abundance varies through the cell cycle, and the nuclear stain reports DNA content, so the cell-cycle position of each cell can be inferred from the same measurements without synchronization.

Here we used the AVITI24 platform to measure transcripts, proteins, phosphoproteins, and cell morphology in the same cells in situ in four human cancer cell lines of lung, colorectal, liver and breast origin. Because this platform is new and has no independent academic track record, we first validated single-cell target detection across cell types and states. We then exposed cells to H_2_O_2_ over a 30-min time course, treated A549 cells to four additional stressors, and co-cultured different cell lines together in the same wells. Within 15 min, chemically distinct stressors changed the same transcripts and phosphoproteins in the same direction and differed mainly in the magnitude of these changes. By 30 min, H_2_O_2_ induced immediate-early genes, most strongly in HCT116 and A549 cells. These changes occurred across the whole population rather than in a responsive subset of cells. The cell-line differences in the immediate-early response persisted in co-culture. MCF7 cells did not respond more strongly when surrounded by strongly responding A549 cells, suggesting that the differences are a property of each cell line rather than of its environment. The response was also similar across inferred cell cycle phases, and the differences among cell lines were larger than the differences between phases. We also tested what the nuclear and whole-cell compartments and a single-cell classifier can and cannot resolve. In situ multiomic measurement within the same cells shows that, during early oxidative stress, a cell’s identity matters more than its neighbors or its place in the cell cycle.

## Results

### Single-cell multiomic measurement of acute stress in four cell lines

To compare the acute stress response among cell types, we profiled lung (A549), colorectal (HCT116), liver (HepG2) and breast (MCF7) cancer cell lines with the AVITI24 Teton MAPK/Cell Cycle panel (**Fig. 1A,B**). The panel measures 338 transcripts (plus GAPDH control probes in each of six imaging batches), 48 protein epitopes (23 phospho-specific) and six cell paint stains in each cell, each quantified in the whole-cell and nuclear compartments (796 features per cell; **Supplemental Table 1**). We refer to each transcript, protein, or stain as a target, and to its measurement in a cellular compartment as a feature. Every feature was detected in every well (Methods). In three runs, we exposed cells to 300 µM H_2_O_2_ for 5, 15 or 30 min alongside untreated controls. In two further runs, we exposed A549 cells to H_2_O_2_ and to four additional stressors. Across the five runs we profiled a total of 1,177,281 cells, of which 1,086,370 (92.3%) passed quality control (**Supplemental Fig. 1A,B**; **Supplemental Table 1**).

**Figure 1.**
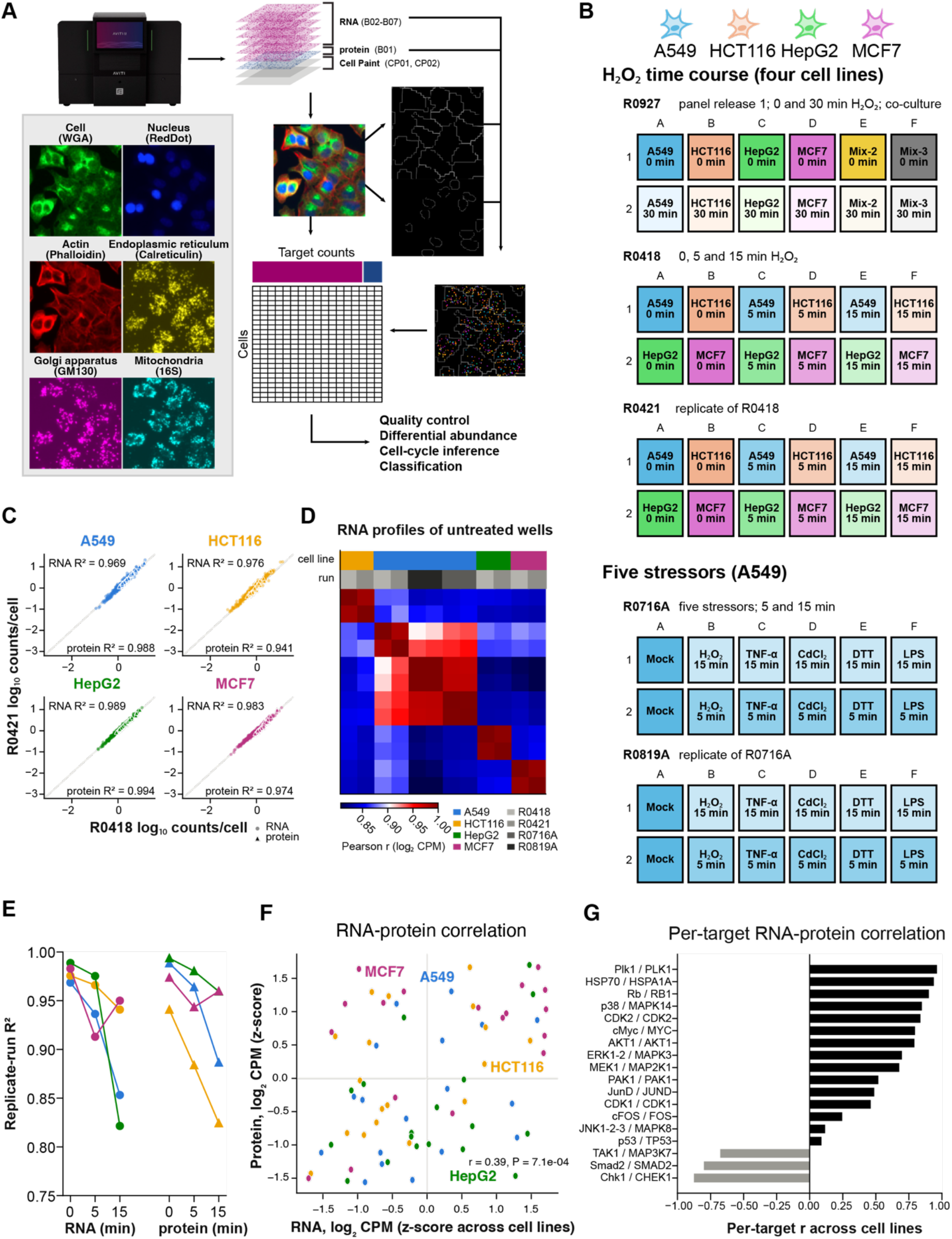
Experimental design, technical reproducibility and multimodal concordance. (A) AVITI24 workflow: cell paint, RNA and protein imaging, segmentation and per-cell target feature extraction. (B) Well layouts of the five runs: three H_2_O_2_ time-course runs with four cell lines and co-cultures, and two runs of A549 cells exposed to five stressors. (C) Per-well mean counts per cell for RNA (circles) and protein (triangles) targets in untreated cells of the two replicate runs. (D) Pearson correlation between RNA profiles (log_2_ counts per million) of all untreated wells from the four panel-v2 runs, hierarchically clustered. Bars indicate cell line and run. (E) Replicate-run R² for every cell line and time point. (F) RNA-protein concordance for the 18 targets measured by both a probe set and an antibody: z-scores of log_2_ counts per million across cell lines. (G) Per-target RNA-protein correlation across the four cell lines for all 18 targets.

Before comparing treatments, we asked whether differences between conditions could be attributed to biology rather than to the run in which the cells were processed (i.e., technical variability). Per-well mean target abundance was highly reproducible between the two replicate runs of the H_2_O_2_ time course (**Fig. 1C**). For untreated cells, R² was 0.97-0.99 for RNA and 0.94-0.99 for protein (**Supplemental Table 1**). If run-to-run variation were large relative to the differences between cell lines, we would expect that the average RNA profiles within each well would group by run. Instead, the average RNA profiles of untreated wells from four runs, performed on two instruments across two chemistry periods, clustered by cell line rather than by run (**Fig. 1D**). Wells of the same cell line correlated more closely, within and across runs (Pearson r = 0.91-1.00), than wells of different cell lines in the same run (r = 0.83-0.88; **Supplemental Table 1**). Reproducibility between the replicate runs was maintained after treatment, with R² above 0.82 for every cell line and time point (**Fig. 1E**). The lower R² of treated wells reflects reduced counts per cell in the 15-min wells of one run and run-to-run differences in response magnitude. The first run used an earlier version of the panel. After harmonization of the two releases by target name (Methods), the run-centered profile of each well in this run correlated best with the matching cell line in the later runs (r = 0.75-0.88; **Supplemental Table 1**). Run-to-run variation was small relative to the differences between cell lines, and therefore data from the two panel releases were analyzed together.

We next validated the *in situ* RNA measurements against an independent method by comparing them with bulk RNA-seq of the same cell lines from the Human Protein Atlas [28, 29]. Across the 338 panel transcripts, AVITI24 abundance correlated with bulk expression within each cell line (Spearman ρ = 0.61-0.72; **Supplemental Fig. 2A**). The differences between cell lines agreed (pooled r = 0.62; median per-gene r = 0.80), and the per-gene correlation was positive for 88% of the 330 genes that varied among the four lines (**Supplemental Fig. 2B**; **Supplemental Table 1**). Consistent with this agreement, the panel profile alone identified each cell line as its own best match among all 1,206 lines in the atlas (**Supplemental Fig. 2C**). The *in situ* RNA measurements therefore reproduce the known expression differences among these cell lines.

RNA and protein levels of the same gene are not necessarily correlated as their synthesis and turnover are regulated separately. However, since both were measured in the same cells, we asked how closely they agreed. Eighteen genes were measured both as transcript and as protein. For 15 of these, relative abundance across the four cell lines was positively correlated between transcript and protein (pooled r = 0.39, P = 7 × 10⁻⁴; **Fig. 1F**). This moderate agreement is consistent with the mRNA-protein concordance reported in bulk and single-cell studies [30, 31]. Agreement was closest for PLK1 and HSP70 (HSPA1A) (r = 0.94-0.96; **Fig. 1G**). Transcript and protein were anticorrelated for TAK1, Chk1 and SMAD2 (r = −0.68, −0.88 and −0.80). Chk1 and SMAD2 protein levels are controlled by ubiquitin-mediated degradation, which may explain why they do not track transcript abundance [32, 33]. Together, these comparisons show that the *in situ* measurements are reproducible across runs, agree with an independent bulk method, and capture RNA and protein information that can be distinct from each other.

### A cell type-invariant early and late response to acute oxidative stress

We next asked which gene products respond to acute oxidative stress (H_2_O2). We summed the RNA and protein counts across all cells within each well and compared each stress time point with untreated cells from the same run using DESeq2 [34]. Testing summed counts per well controls false discoveries better than treating individual cells as replicates [34–36]. DESeq2 normalizes each well by a size factor, the median ratio of its counts to those of the other wells, which corrects for differences in probe yield between wells. For example, the 15-min treated A549 and HepG2 wells of one run had about 35% fewer RNA counts per cell than the untreated wells (**Supplemental Fig. 1C**). This normalization assumes that most targets do not change. This held here, because at most a third of the transcripts changed in any single comparison (Methods). In total, 93 targets (79 transcripts and 14 proteins) changed significantly (FDR < 0.05) in at least one cell line and time point (**Fig. 2A**; **Supplemental Table 2**). The number of responsive targets peaked at 15 min in every cell line (A549, 41; MCF7, 47; HCT116, 30; HepG2, 12; Fig. 2B). Two additional A549 runs, performed for the stressor comparison below, each contained H_2_O_2_-treated wells at 5 and 15 min and mock-treated wells. We compared each H_2_O_2_ well with the mock wells of the same run and analyzed these runs together with the two time-course runs. All 41 targets that responded at 15 min in the time-course runs changed in the same direction in this four-run analysis, 33 of them significantly, and fold changes in the two analyses were correlated (r = 0.92; **Supplemental Table 2**). All four cell lines responded to H_2_O_2_ and the A549 response was reproduced in independent runs.

**Figure 2.**
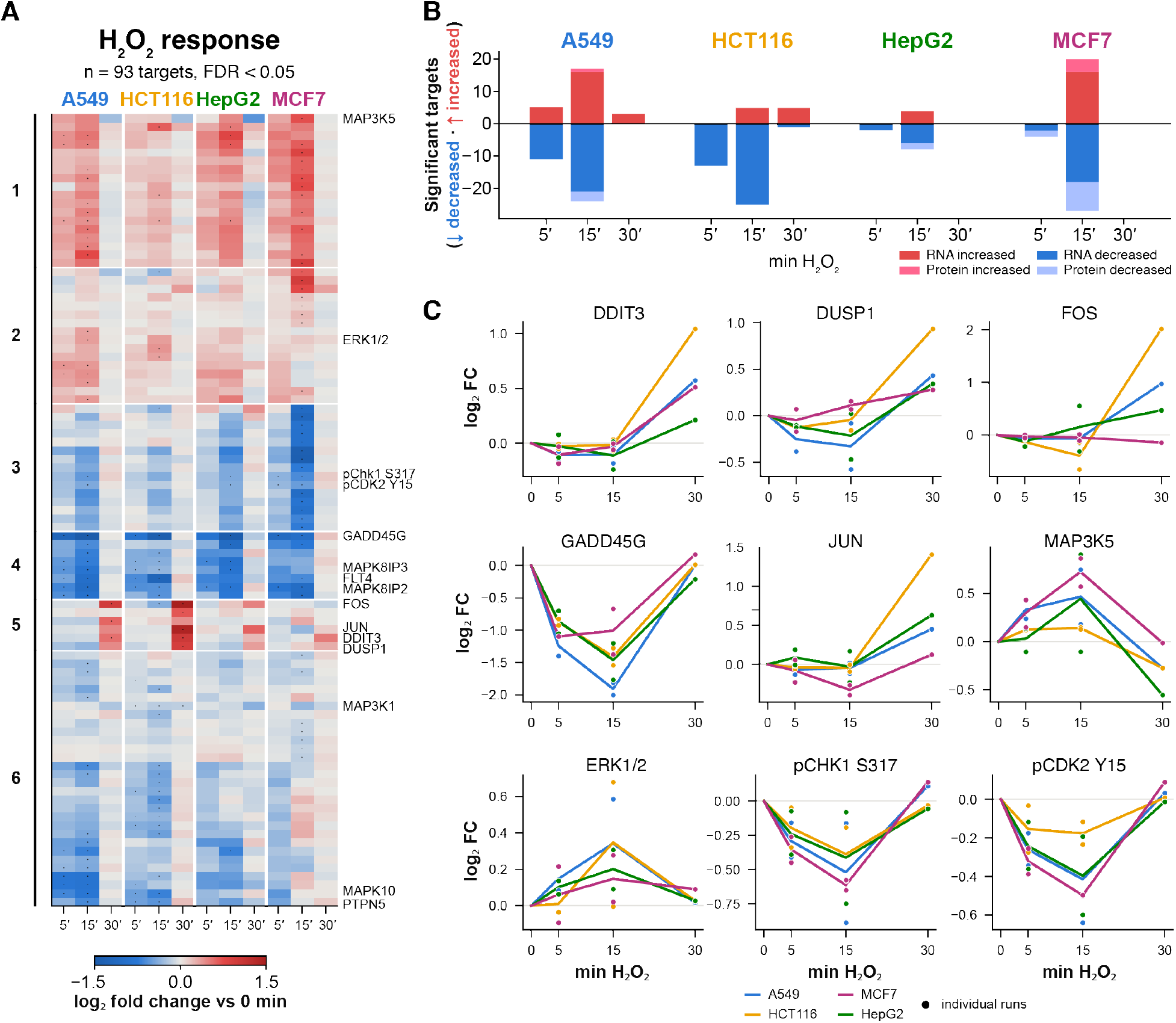
Replicate-aware response to 300 µM H_2_O_2_. (A) log_2_ fold change versus untreated cells of the same run (PyDESeq2, ∼ run + time) for the 93 targets significant (FDR < 0.05) in at least one cell line and time point. Dots mark significant cells of the heatmap, and rows are clustered (Ward) into six groups. (B) Number of significantly increased and decreased targets per cell line and time point. Lighter segments are proteins. (C) Example targets: lines show model estimates and points show individual runs (size-factor-normalized). The 30-min estimates derive from a single run.

Measuring signaling proteins and transcripts in the same cells allowed us to resolve how the response is organized in time. The responsive targets fell into two groups (**Fig. 2A,C**). Most (87 of 93) changed within 5-15 min and returned to near untreated levels by 30 min. This rapid layer was dominated by losses (53 of 87), led by a decrease in GADD45G transcripts in all four cell lines, together with lower phosphorylation of CHK1 and CDK2 at checkpoint-associated sites. The remaining six targets were induced only at 30 min. Five of them, FOS, JUN, DUSP1, DUSP2 and NR4A1, are classical immediate-early genes, the first genes that cells transcribe after a stress. The sixth, DDIT3 (CHOP), is a stress-induced transcription factor. All six are induced by oxidative stress in bulk studies [19, 37]. In all four cell lines, we found that H_2_O_2_ first triggers a rapid signaling layer of phosphorylation changes and transcript losses within minutes, and then induces new transcription of immediate-early genes by 30 min.

Although the order and direction of the response were shared across cell lines, the magnitude of response was not. Induction of the immediate-early genes ranged from a mean log_2_ fold change of 1.1 in HCT116 cells and 0.5 in A549 cells to 0.3 in HepG2 and 0.1 in MCF7 cells. It was significant in HCT116 (*FOS* log_2_ fold change 2.0, FDR = 4 × 10⁻²⁷) and A549, but not in HepG2 and MCF7. HepG2 also showed the fewest changes at 5 and 15 min (**Fig. 2A,B**). Therefore, cell lines share the direction of the response to H_2_O_2_ but differ in how strongly they respond to oxidative stress.

Bulk measurements cannot distinguish a small response in every cell from a larger response in a subset of cells. To distinguish these possibilities, we compared the per-cell count distributions of eight example targets before and after treatment (**Supplemental Fig. 3**). We measured cell-to-cell variability as the Fano factor, the variance of per-cell counts divided by their mean [38]. For the same increase in mean expression, a response confined to a subset of cells would increase the Fano factor much more than a uniform shift of the whole population. We examined the 12 cases in which an example target increased at least 1.4-fold on average, 11 of them immediate-early genes at 30 min. In these cases, the Fano factor increased by a median of 1.19-fold. This increase was smaller than expected for a uniform shift of all cells (1.37-fold, given the baseline variability of these counts) and well below the 1.7-and 2.5-fold increases expected if only half or one fifth of the cells had responded (**Supplemental Table 2**). This is consistent with induction through more frequent transcription in nearly all cells rather than activation of a responding subpopulation. Together, these results show that H_2_O_2_ induces a two-stage response that is shared in direction by all four cell lines, differs among them in magnitude, and occurs across the whole cell population.

### Chemically distinct stressors elicit a common early response in A54G

To determine whether the early response is specific to oxidative stress, we exposed A549 cells to four other stressors that act through distinct mechanisms: the inflammatory cytokine TNF-α, the heavy-metal cadmium, the reducing agent dithiothreitol (DTT) and the bacterial outer-membrane component lipopolysaccharide (LPS). Cells were exposed for 5 and 15 min in two independent runs, and each condition was compared with mock-treated cells of the same run (**Fig. 3**; **Supplemental Table 2**). Each condition significantly altered the expression of 112-263 probed RNA and protein targets (FDR < 0.05; **Fig. 3A**). Most changes were shared across stressors, supporting the concept of a common environmental response.

**Figure 3.**
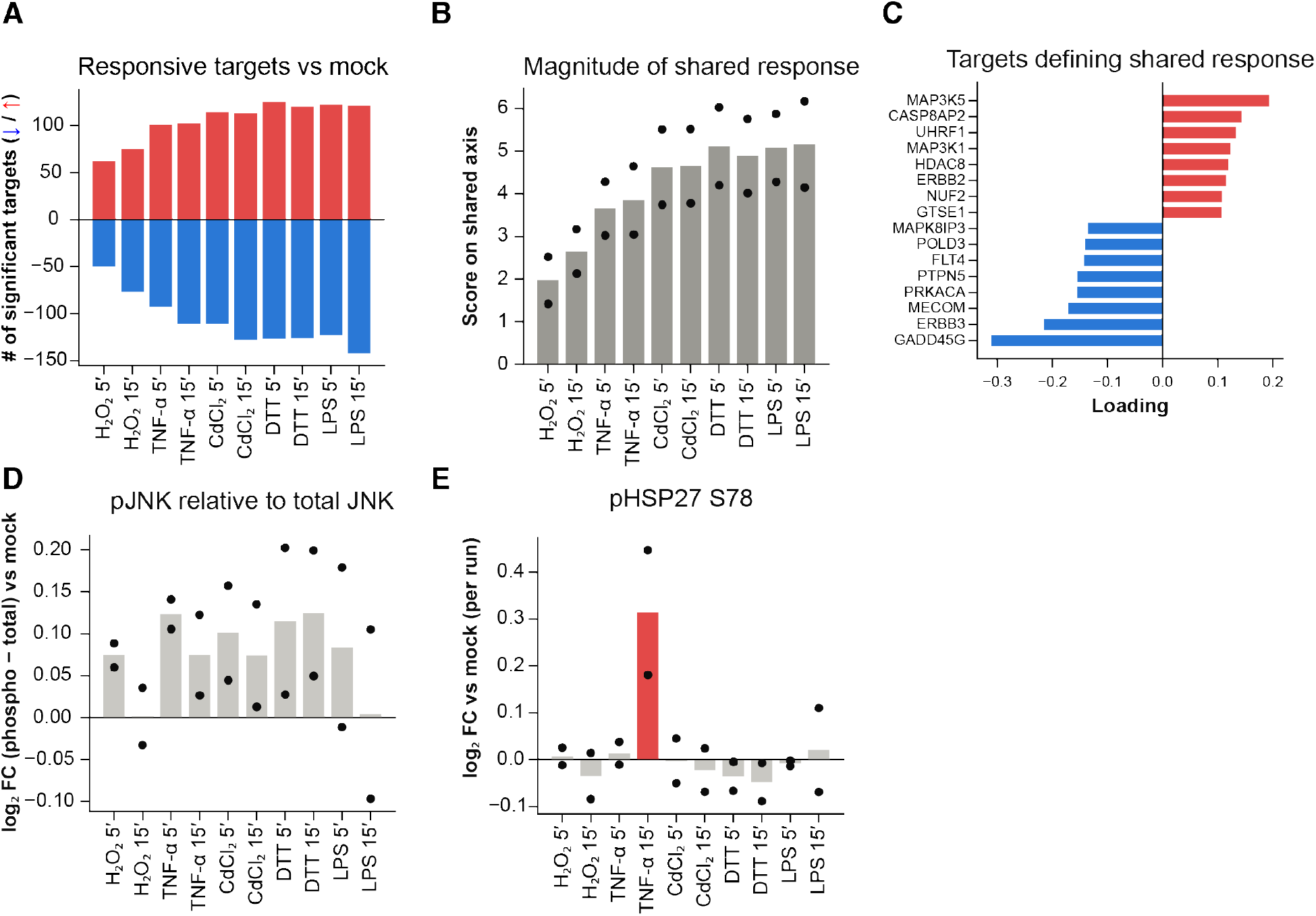
Chemically distinct stressors produce a dominant shared response profile. A549 cells were exposed to H_2_O_2_, TNF-α, CdCl_2_, DTT or LPS for 5 or 15 min (two runs) and compared with mock-treated cells of the same run (PyDESeq2, ∼ run + condition). (A) Number of significantly increased (red) and decreased (blue) targets (FDR < 0.05). (B) Magnitude of each condition along the first singular vector of the fold-change matrix of responsive targets (98.6% of the squared fold-change magnitude). Bars are means and points are runs. (C) Targets with the largest loadings on the shared axis. (D) Phospho-JNK1/2/3 relative to total JNK1/2/3 (difference of log_2_ fold changes versus mock) per run. (E) Phospho-HSP27 Ser78, log_2_ fold change versus mock per run.

If the stressors acted through one shared program, their fold changes should differ only by a scaling factor. Across the 321 targets in A549 that responded to at least one stressor, the fold changes of every pair of conditions were highly correlated (r ≥ 0.94). To measure the size of each response, we decomposed the target × condition matrix of fold changes by singular value decomposition. The first component, which represents the shared profile, accounted for 98.6% of the squared fold-change magnitude (**Fig. 3B**). The score of each condition on this component measures the size of its response. H_2_O_2_ produced the smallest response, followed by TNF-α and cadmium, and DTT and LPS produced the largest. Except for H_2_O_2_, each stressor’s response was already as large at 5 min as at 15 min. The five stressors therefore engage the same early program and differ mainly in how strongly they engage it.

The targets that contributed most to the shared profile were those that changed early in the H_2_O_2_ time course, including decreases in GADD45G, FLT4, PRKACA and MAPK8IP3 and increases in MAP3K5 and MAP3K1 (**Fig. 3C**). Several of these encode components of the JNK pathway, which suggests that JNK activation might set the size of the shared response. If so, JNK phosphorylation should rise with the magnitude of each stressor’s response. Phosphorylation of JNK relative to total JNK increased modestly in 17 of the 20 stressor-treated wells (mean +0.08 log_2_; **Fig. 3D**), but it did not rise with the magnitude of the shared response (r = −0.38 across wells). JNK was therefore modestly activated in nearly all stressor-treated wells, but the level of its activation was not predictive of how strongly each stressor engages the shared program.

We next asked whether any targets responded specifically to a given stressor. The panel does not include transcriptional readouts that typically distinguish these stressors, such as NRF2 targets for oxidants (HMOX1, NǪO1), NF-κB targets for TNF-α and LPS (NFKBIA, TNFAIP3), metallothioneins for cadmium, or unfolded-protein-response genes for DTT (HSPA5, XBP1). Our analysis can only detect stressor-specific changes among the measured MAPK and cell-cycle targets. After removal of the shared component, the remaining stressor-specific variation was small (residual s.d. 0.03 versus 0.24 log_2_ units for the fold changes), with one exception. Phosphorylation of HSP27 at Ser78 increased after 15 min of TNF-α in both runs (log_2_ fold change 0.30, FDR < 10⁻⁴) and in no other condition (**Fig. 3E**), consistent with activation of p38-MAPKAPK2 downstream of the TNF receptor [20, 39, 40]. Among the measured targets, the five stressors share a single early response, and TNF-α-induced HSP27 phosphorylation is the only stressor-specific event detected.

From this analysis we conclude that within the first 15 min, the MAPK and cell-cycle regulators measured by the panel respond to chemically distinct stresses through a common program whose amplitude depends on the stressor. Pathway-specific events, such as TNF-α-induced HSP27 phosphorylation, occur in addition to this program. Stressor-specific programs may emerge at later times or in genes outside the panel.

### The stress response is not detectably transferred to neighboring cells

Specific stresses typically affect some organs more than others. For example, the liver removes environmental toxins, and the skin is the first line of exposure to UV light. Cells from different tissues may therefore be tuned to respond to particular stresses. The four cell lines did differ in how strongly they induced the immediate-early genes at 30 min, from strongest in HCT116 cells to weakest in MCF7 cells (**Fig. 2A,C**). Because each cell line was grown in its own well, however, these differences could also reflect the well rather than the cell line. Cells remove extracellular H_2_O_2_ at a rate proportional to their number [41, 42], so the effective dose depends on how many cells a well contains, and the four lines grew to very different densities (**Supplemental Fig. 1A**). Each condition also occupied a single well per run. Growing the cell lines together in the same well equalizes medium, dose, cell density and handling, and therefore tests whether the differences are intrinsic to each line.

Co-culturing distinct cell lines can also test whether a stress response spreads between cells. If the detected differences in oxidative stress response arose as a function of homogenous growth in dedicated wells, the differences should disappear when the cell lines share a well. Additionally, if a strongly responding cell line transmitted its response to its neighbors, weakly responding cells should respond more strongly in co-culture. We therefore grew A549 and MCF7 cells together (Mix-2), and A549, MCF7 and HepG2 cells together (Mix-3), alongside pure cultures of each line (**Fig. 4A**; **Supplemental Fig. 4A**). All cells were then exposed to H_2_O_2_ for 30 min or left untreated. We assigned each cell to a cell line with a classifier trained on the pure cultures (99.4-100% accuracy on held-out pure-well cells; **Supplemental Table 3**). The immediate-early probes were removed from this classifier so that cell-line assignments would not depend on the response being measured. To compare the immediate-early response of different cell lines in the same well, we defined an immediate-early gene (IEG) score for each cell. The score is the number of FOS, JUN, DDIT3, DUSP1 and NR4A1 counts per 10⁴ RNA counts of the cell. These are the genes induced at 30 min (**Fig. 2A**), except DUSP2, which was induced only in HCT116 cells, a cell line not included in the co-cultures. Normalizing to each cell’s total RNA counts corrects for differences in total detected RNA among cells and cell lines.

**Figure 4.**
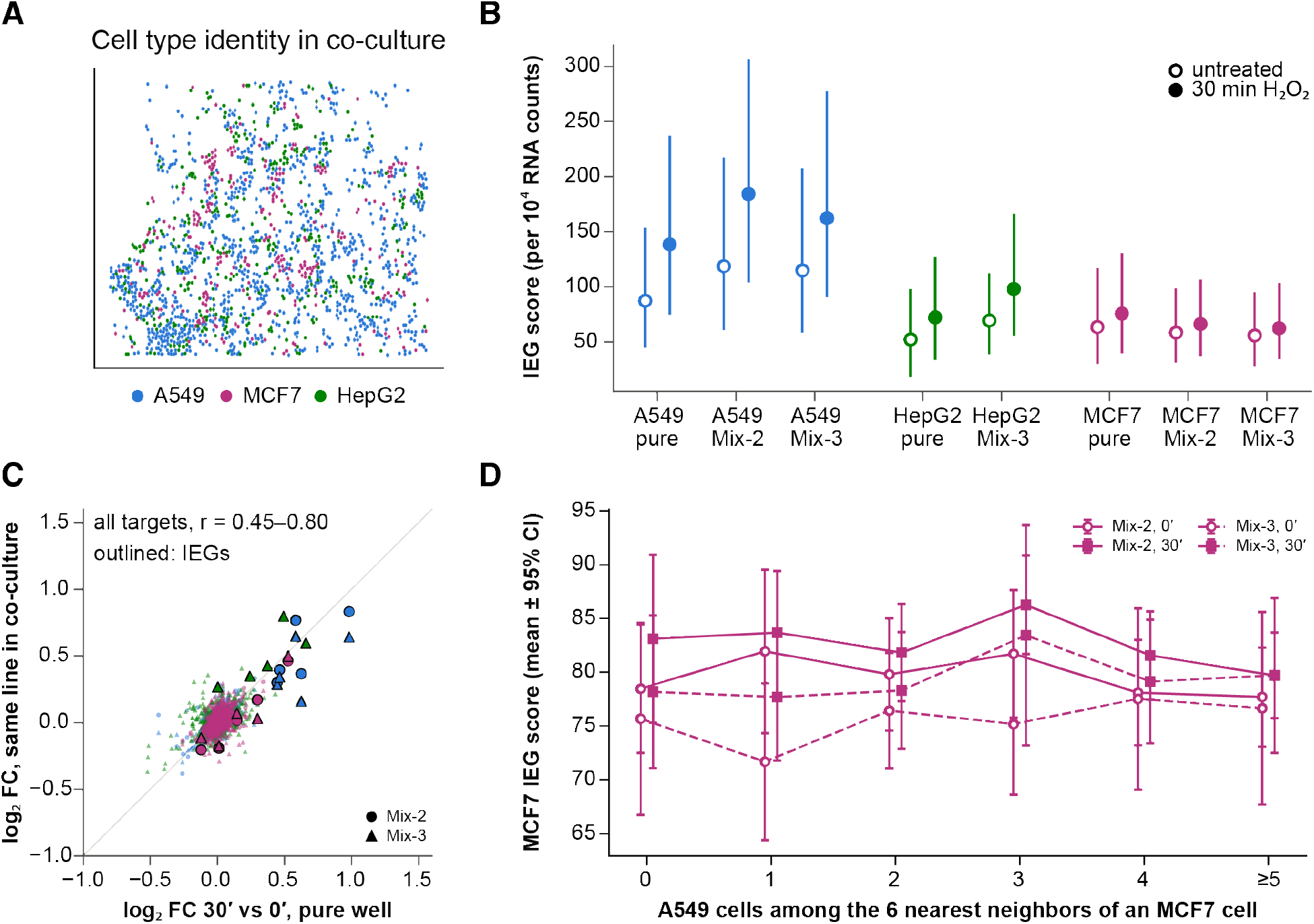
Cell-line differences in the immediate-early response persist in co-culture without detectable local transfer. (A) Positions of cells in one imaging tile of a co-culture of A549, MCF7 and HepG2 cells, colored by predicted cell line (classifier trained on pure wells without the immediate-early probes). (B) IEG score (median and interquartile range) of each cell line in its pure well and in co-culture, untreated and after 30 min of H_2_O_2_. (C) Fold changes (30 vs 0 min) of all targets in co-culture versus the pure well. IEGs are outlined. (D) IEG score of MCF7 cells in co-culture versus the number of A549 cells among their six nearest neighbors (mean ± 95% CI), untreated and after 30 min of H_2_O_2_.

In co-culture, the median IEG score of the strongly responding A549 and HepG2 cells increased after 30 min of H_2_O_2_ (by 41-55% and 41%, respectively), whereas that of MCF7 cells increased only weakly (by 11-13%), as in pure culture (**Fig. 4B**; **Supplemental Fig. 4B**). The difference between the cell lines persisted when they shared a well, indicating differences do not arise from differences between wells. Co-culture did raise the baseline expression of some of these genes in A549 cells. Untreated A549 cells in co-culture expressed *FOS*, *JUN*, *DUSP1* and *NR4A1* at 0.3-0.9 log_2_ units higher than in the pure well, whereas *DDIT3* was 0.2-0.3 units lower (**Supplemental Table 3**). Neighboring cells can therefore influence the baseline expression of some genes, although these effects were small.

The agreement between mixed and pure wells extended beyond the immediate-early genes. The 30-min fold changes of all targets in co-culture correlated with those in the pure wells (r = 0.45-0.80; **Fig. 4C**). For A549 and MCF7, the fold changes in mixed and pure wells differed no more than the fold changes in the two co-cultures differed from each other (s.d. of fold-change differences 0.06-0.08 versus 0.06-0.07 log_2_ units; HepG2, present only in Mix-3, 0.12; **Supplemental Table 3**). Apart from NR4A1 and HSP60 (both in A549), sharing a well did not change how each line responded to H_2_O_2_. The response program of each cell line is therefore an intrinsic property that is not measurably influenced by other cell types in the same well. This comparison averages over all cells of a line, however, and could miss an effect confined to cells that are adjacent to another cell line.

Because the position of every cell is recorded, we could test for local effects directly. If the immediate-early response were transmitted through contact-dependent or short-range signals, MCF7 cells should respond more when more of their neighbors are strongly responding A549 cells. We did not detect this association. After 30 min of H_2_O_2_, the IEG score of MCF7 cells did not change with the number of A549 cells among their six nearest neighbors (slope −0.7 and +0.5 per neighbor, P = 0.27 and 0.61), as in untreated wells (**Fig. 4D**).

To determine how large an undetected effect could be, we modeled the stress-induced change in this relationship (Methods). The upper 95% confidence bound on this neighbor effect was 2-4% of the difference between the A549 and MCF7 responses per adjacent A549 cell (5-7% against a permutation null that shuffled cell identities within each imaging tile). None of ten neighborhood definitions, including fixed radii and distance-weighted exposure, detected a positive effect (**Supplemental Fig. 4C**; **Supplemental Table 3**). Within 30 min and within the resolution of this assay, the immediate-early response is not transmitted from strongly responding cells to their weaker neighbors.

The co-cultures also allowed us to measure two technical properties of the platform. Because each cell’s identity and position are read out *in situ*, the composition of a mixed population, including the identify of cell neighbors, can be measured directly rather than inferred from seeding ratios. Measured composition was reproducible across replicates but departed from the seeding plan. A549 cells were over-represented 1.3-1.5-fold relative to its expected seeding ratio (**Supplemental Fig. 4D**), possibly because of differences in attachment or proliferation, and cells clustered modestly with cells of the same type (**Supplemental Fig. 4E**). Adjacent cells of different types also express different markers, which allowed us to estimate how much signal is assigned to the wrong cell, a known source of error on other *in situ* platforms [43]. Each adjacent cell of a different type contributed marker signal equal to 0.8-0.9% of that cell’s own level (**Supplemental Fig. 4F**). Some of this signal may reflect low-level expression of the marker in the receiving cell, so this value is an upper estimate of misassignment. At this level, adjacent cells of different types can be analyzed as separate populations.

Cell positions also allowed us to ask whether local cell density affects the stress response (tested in homogeneous cultures). Cells in crowded regions may have a greater collective ability to detoxify, and so might be exposed to less H_2_O_2_ per cell, resulting in a diminished response [41, 42]. This was the case in HCT116 cells, which grow in compact colonies and reached the highest local densities. At 15 min, seven early-response transcripts decreased less in the most crowded fifth of cells in each well than in the sparsest fifth, a 20-26% weaker response that was consistent in both time-course runs (**Supplemental Fig. 5A**). The immediate-early induction at 30 min was also weaker in crowded cells (log_2_ fold change 0.84 versus 1.24; **Supplemental Fig. 5B**).

The difference in early-gene response remained significant after adjustment for cell size, total RNA and cell-cycle position, in both runs and at both 5 and 15 min. The adjusted immediate-early difference had P = 0.03 (0.39 without adjustment for total RNA). MCF7 cells showed the same trend, whereas the responses of A549 and HepG2 cells did not depend on local density (**Supplemental Fig. 5C**; **Supplemental Table 3**). While we did not measure local H_2_O_2_, local density nevertheless dampened the response in HCT116 cells and should be considered when comparing cell types or conditions that differ in crowding. Together, the co-culture and density analyses show that cell-line differences in the acute response are intrinsic to each line. Neighboring cells may still influence one another indirectly, by lowering each other’s exposure through detoxification, rather than by transmitting the response. These experiments used artificial mixtures of cell lines from different tissues, and whether neighbors influence the stress response in natural tissue remains an open question.

### Multiomic profiling identifies where each cell is in the cell cycle

Cell-cycle state is a major source of variation among proliferating cells, and bulk studies usually control it by chemical synchronization, which itself induces a stress response [16, 17]. To avoid synchronization, we sought to infer the cell-cycle position of each cell from cell cycle marker expression profiles. Our panel of RNA probes includes 40 of the 43 S-phase genes and 53 of the 54 G2/M genes commonly used for single-cell cell-cycle scoring [25]. We scored each cell for the S and G2/M programs and assigned cells with low scores for both to G1 as previously described [25]. We ordered cells using only these 93 reference genes, leaving other panel targets that may also vary with the cycle for independent comparison.

S-phase genes are expressed before G2/M genes, and both fall after division, when cells re-enter G1. Cells progressing through the cycle should therefore fall in a consistent order in a projection of these genes onto their first two principal components, computed from untreated cells (Methods). In all four cell lines, G1, S and G2/M cells fell in the same order around the center of this projection (**Fig. 5A**). We assigned each cell a cell-cycle position (0 to 1) around a circular plot (angle around this center), starting at G1 (Methods). As expected from the genes used to construct this ordering, the S score peaked first (position ≈ 0.3) and the G2/M score second (≈ 0.7) in all four cell lines (**Fig. 5B**). As an independent test, we analyzed DNA content from nuclear DNA staining produced by AVITI24 imaging, which was not used to construct the ordering. DNA content rose from 1.0 to 1.5-1.8 times the level observed in G1, between positions 0.4 and 0.7, and returned to 1.0 after the G2/M peak (**Fig. 5C**), the profile expected for DNA replication followed by cell division.

**Figure 5.**
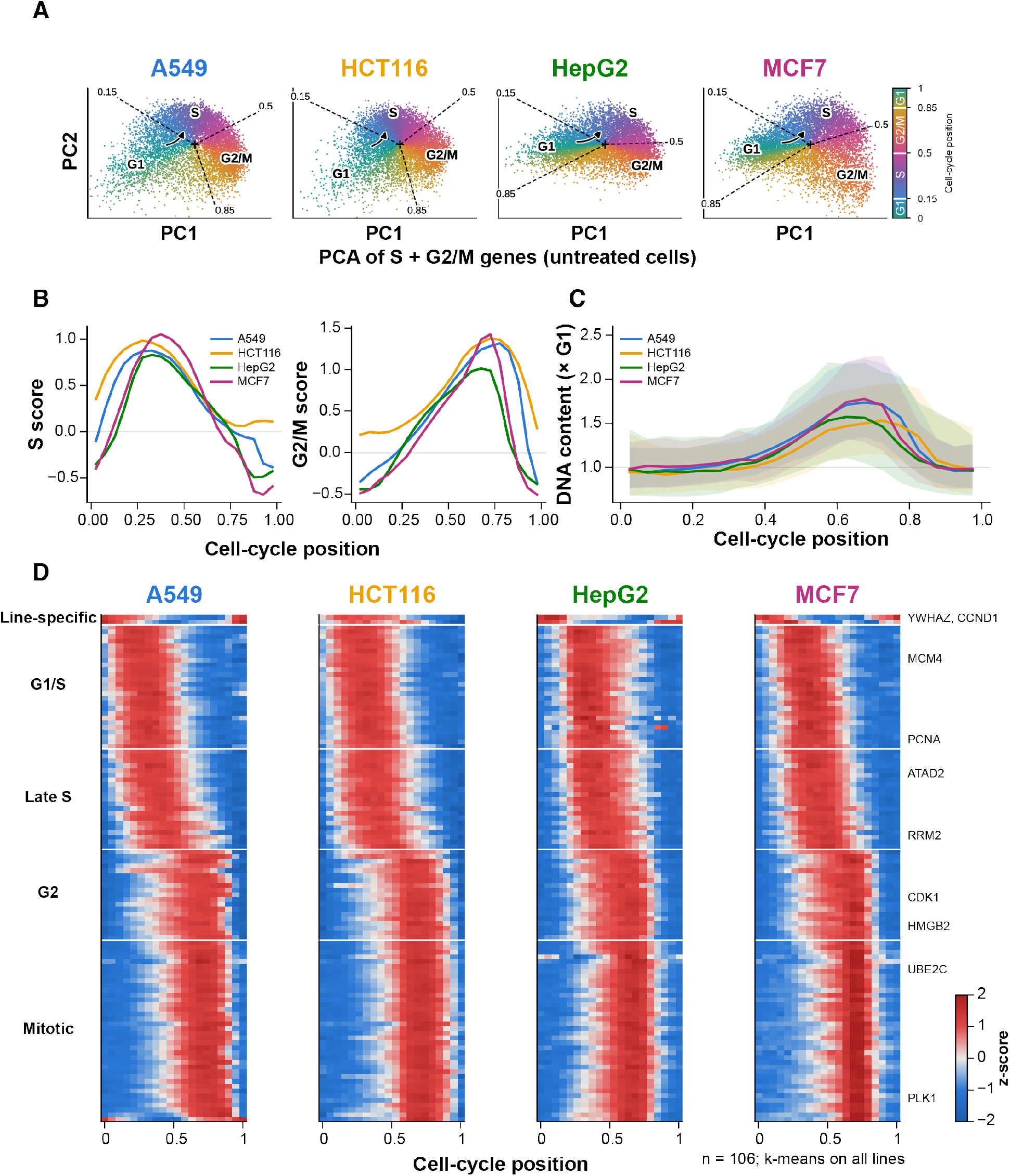
Cell-cycle position inferred from the multiomic profile. (A) Projection of RNA expression in untreated cells onto the first two principal components of 93 S and G2/M genes, colored by cell-cycle position (angle around the centroid +, anchored at G1 and rank-transformed). Dashed lines mark the cell cycle phase boundaries (positions 0.15, 0.5 and 0.85), and the arrow indicates the direction of progression. (B) Mean S and G2/M scores along the cell-cycle position. (C) DNA content (integrated nuclear DNA-stain signal relative to G1 cells of the same well; median and interquartile range) along the cell-cycle position. (D) Mean expression of the 106 cycle-associated transcripts in 20 position bins (z-scored within each cell line), clustered by k-means on the concatenated profiles of all four cell lines and ordered by peak position.

We next identified the panel targets whose abundance changes along this ordering. In total, 106 transcripts varied with inferred cell-cycle position (**Fig. 5D; Supplemental Table 4**). Of these, 86 are reference genes used to build the ordering, so they vary along it partly by construction. The remaining 20 were not used, and they peaked at the stages expected from their known functions, for example MCM3 and CDT1 at G1/S and PLK1 and CCNB1 in mitosis. Clustering by k-means separated the transcripts into four groups that peaked in sequence, at G1/S, late S, G2 and mitosis (**Fig. 5D**). A fifth, small group (CCND1 and YWHAZ) varied with the cycle in only two of the four cell lines. The profiles were similar across the four cell lines (median correlation between lines r = 0.92), even though expression levels differed. No protein passed the threshold for cycle dependence. The most cycle-dependent protein feature, CDK2 Tyr15 phosphorylation, explained at most 3% of variance and peaked at G2/M in all four lines. Together, the score profiles, the DNA content and the 20 genes not used for the ordering show that the inferred position follows progression through the cycle in the same way in all four cell lines. The panel alone is therefore sufficient to order cells along the cycle and to compare cell-cycle positions across cell lines without synchronization.

### The acute stress response is largely independent of the cell cycle

We next asked whether the response to H_2_O_2_ depends on cell-cycle phase. Because 30 min is much shorter than the cell cycle (typically >20 hrs), H_2_O2 did not consistently change the fraction of cells in G1, S, or G2/M (**Fig. 6A**). We could therefore compare the stress response within each cell cycle phase. In doing so, baseline differences between phases, such as the periodic expression of cell-cycle genes (**Fig. 5D**), are removed, and so any remaining difference reflects the response intrinsic to H_2_O_2_. Measured this way, responsive targets changed in the same direction and by similar amounts in G1, S, and G2/M cells (**Fig. 6B,C**). Their fold changes were highly correlated between phases in every cell line and at every time point (r = 0.72-1.00) and were nearly identical in G1 and G2/M cells (r = 0.96; **Fig. 6D**). The same held for the 93 cell-cycle reference genes, 20 of which responded to H_2_O_2_ at 15 min (r = 0.88-0.90 between G1 and G2/M in each cell line), and for early-response targets that themselves regulate the cell cycle: CDK2 Tyr15 phosphorylation, CHK1 Ser317 phosphorylation and GADD45G decreased by similar amounts in every phase. So when cells were grouped into three discrete phases, the response to H_2_O_2_ did not depend on cell-cycle phase.

**Figure 6.**
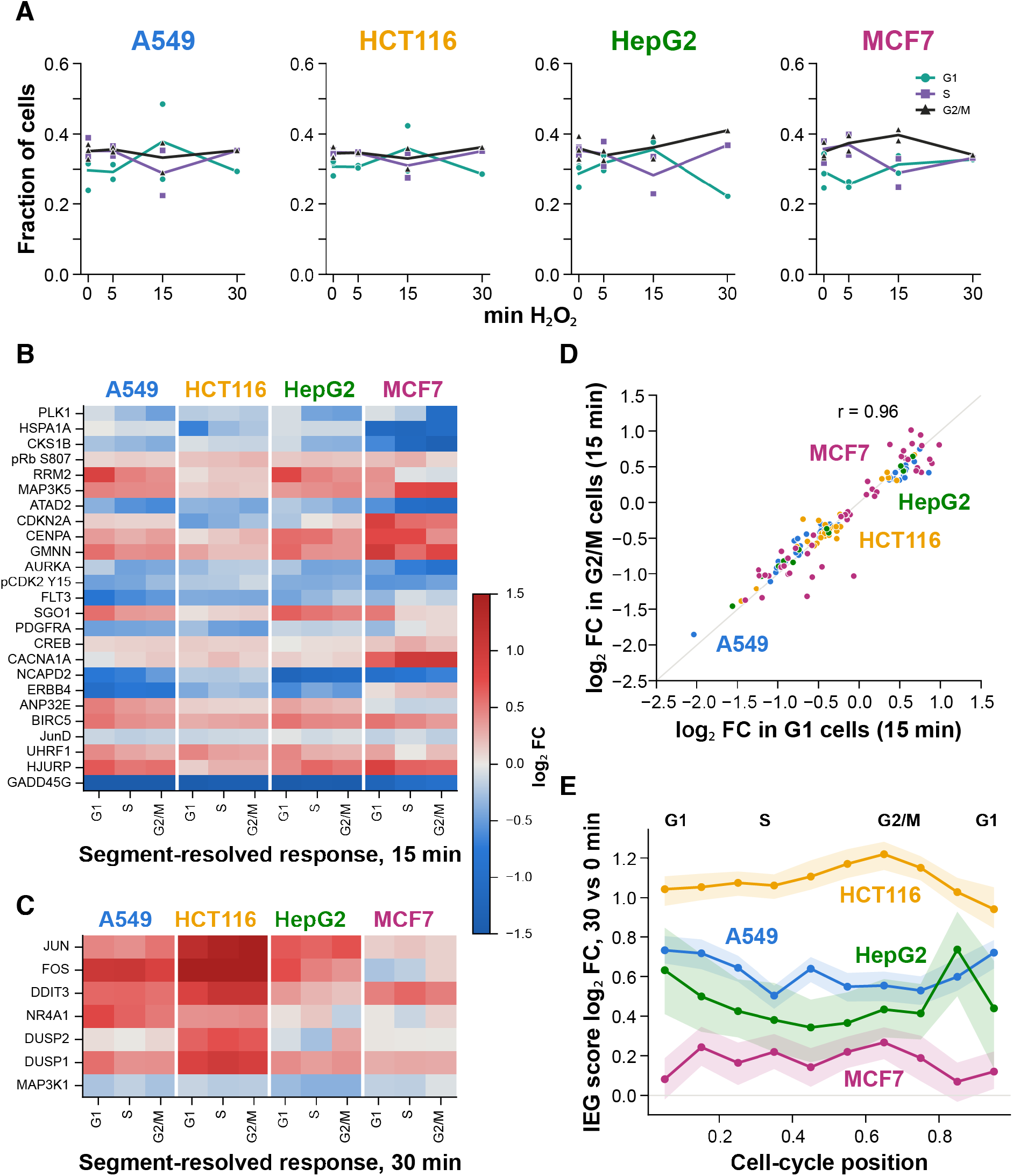
Response variation among cell lines and cell-cycle phases. (A) Fraction of cells in the G1, S and G2/M phases of the cell-cycle position per well (points) and mean across runs (lines). (B,C) Response to H_2_O_2_ within each cell-cycle phase at 15 min (B) and 30 min (C). Each cell shows the log_2_ fold change of a target in treated versus untreated cells of the same phase (PyDESeq2), for each cell line and phase. B shows the 25 of 87 targets responsive at 15 min whose responses differed most between phases. C shows all 7 targets responsive at 30 min. (D) Fold changes in G1 versus G2/M cells at 15 min for responsive targets. (E) Fold change of the immediate-early gene score (30 vs 0 min) in ten cell-cycle position bins for each cell line (bootstrap 95% confidence intervals).

We checked whether the broad cell-cycle grouping masked finer temporal responses, using two more sensitive analyses. Across ten finer cell-cycle bins, the immediate-early induction at 30 min varied by only 0.20-0.28 log_2_ units within each cell line (0.39 in HepG2, whose bins contain fewer cells). It was slightly stronger in G1 in A549 and in S/G2 in HCT116, but these differences were several-fold smaller than those between cell lines (**Fig. 6E**). A comparison of G1 and G2/M cells within the same well, which removes well-to-well variation, detected phase differences for only three responses, all in HCT116 and each 1.4-fold or less (JUN and DDIT3 at 30 min, HSPA1A at 15 min; **Supplemental Table 4**). The cell cycle therefore modulates only a few responses, and only slightly, whereas the magnitude of the acute response to H_2_O_2_ is set mainly by cell identity.

### Subcellular localization of RNA and proteins

Because every target is quantified both within the nucleus and across the whole cell, we next examined how targets partition between these compartments during the stress response. The images are two-dimensional, so the nuclear compartment also contains cytoplasm lying above and below the nucleus. The cell-paint stains show the extent of this overlap. Relative to its share of cell area, DNA staining was enriched 1.8-3.5-fold in the nuclear compartment, but stains for the endoplasmic reticulum, Golgi and mitochondria were also enriched, by 1.1-2.2-fold (**Supplemental Table 5**). Consistent with this, proteins with predominantly nuclear localization (for example Rb, phospho-Rb Ser807, p27, c-Fos, and JunD) were only moderately more enriched than cytoplasmic or mitochondrial proteins (for example HSP60, phospho-PKA, and PAK1; AUC = 0.68, one-sided P = 0.04; **Fig. 7A**). The nuclear compartment therefore cannot definitively assign individual targets to the nucleus, but changes in a target’s nuclear fraction between conditions can indicate redistribution between the nuclear region and the rest of the cell.

**Figure 7.**
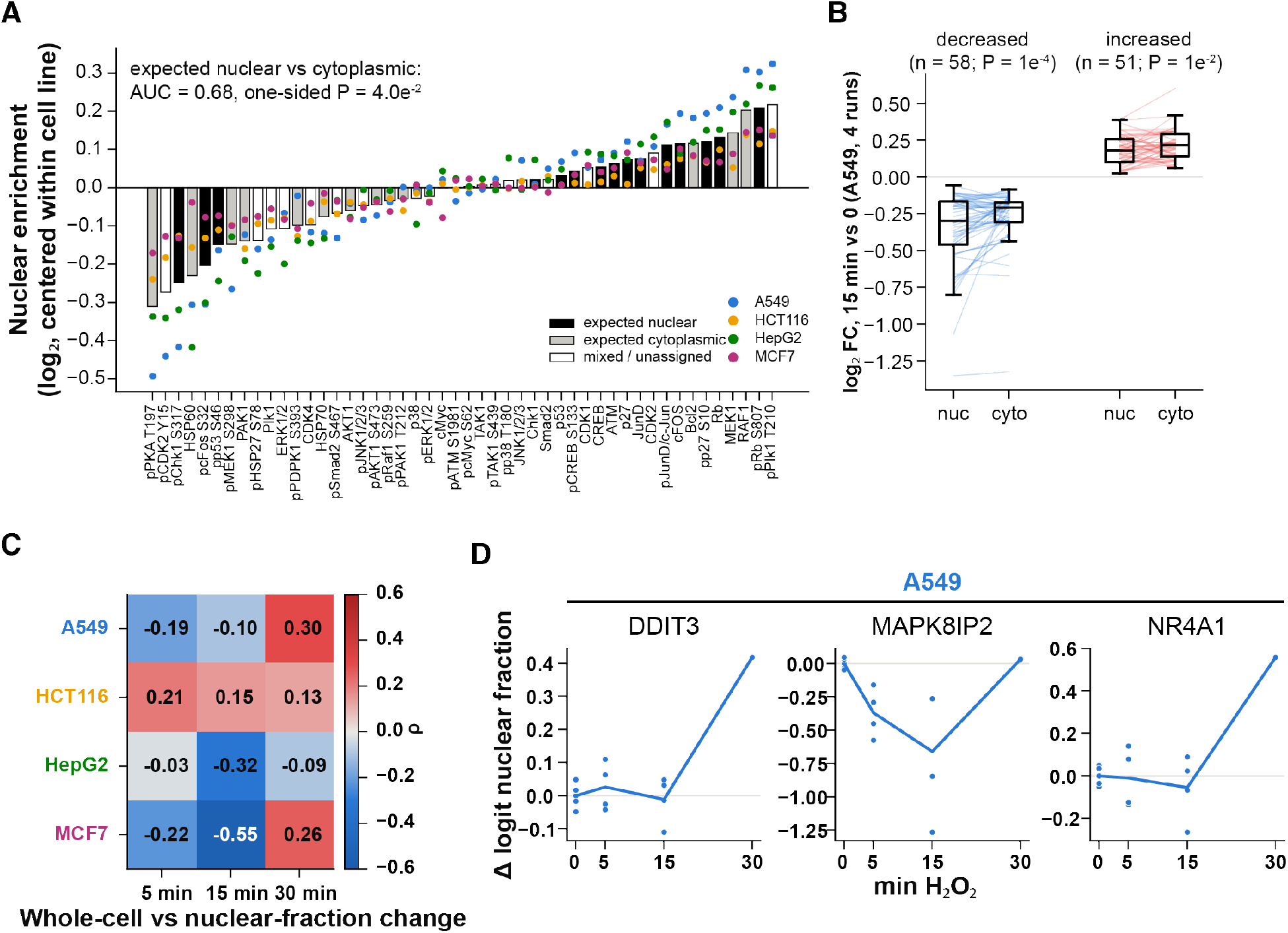
Subcellular partitioning of targets and of the stress response. (A) Nuclear enrichment of each protein in untreated cells (nuclear fraction divided by the nuclear share of cell area; log_2_, centered within each cell line). Bars show the mean across cell lines, colored by literature-expected localization, and points show individual cell lines. (B) Nuclear and cytoplasmic log_2_ fold changes at 15 min of H_2_O_2_ for transcripts significantly decreased or increased in A549 cells (four runs). (C) Spearman correlation between whole-cell log_2_ fold change and change in logit nuclear fraction across transcripts, per cell line and time point. (D) Illustrative examples of nuclear-fraction changes in A549 cells: change in logit nuclear fraction of MAPK8IP2, NR4A1 and DDIT3 transcripts relative to untreated cells of the same run (points are runs; lines are means). These transcripts were chosen from the few with significant changes (FDR < 0.05; Supplemental Table 5) to illustrate the 15-min decrease (MAPK8IP2) and the 30-min increase (NR4A1, DDIT3). They are not representative of all targets, most of which showed no significant change.

We next asked whether transcript changes after H_2_O_2_ were greater in the nucleus, where newly synthesized transcripts first accumulate, or in the cytoplasm. In A549 cells (across four runs), we took the transcripts whose abundance decreased at 15 min and compared the size of the decrease in each compartment. The decrease was larger in the nuclear compartment than in the cytoplasm (median log_2_ fold change −0.30 vs −0.21, Wilcoxon P = 1 × 10⁻⁴; **Fig. 7B**). This is consistent with reduced transcription, which would deplete nuclear transcripts first, or with faster export of transcripts from the nucleus.

Induced transcripts showed the mirror image at 30 min. The immediate-early transcripts that increased in A549 and HCT116 also shifted toward the nucleus, with a larger fraction of their copies located there than in untreated cells (**Fig. 7C,D**; **Supplemental Table 5**), as expected if newly transcribed RNA accumulates in the nucleus before export. For the transcripts that changed most, the change was larger in the nucleus in both directions. Across the panel as a whole, this coupling was weaker at 5 and 15 min than at 30 min. At the early times, the correlation between a transcript’s change in abundance and its change in nuclear fraction was positive in HCT116 but negative in A549, HepG2, and MCF7 (**Fig. 7C**).

Individual nuclear-fraction changes were rarely significant, so we assessed the compartment measurements as a whole rather than by selected examples. Compartment abundance was highly reproducible. Each transcript’s nuclear fraction in untreated cells correlated strongly between the two replicate runs (r = 0.94-0.99), and its change at 15 min correlated moderately (r = 0.40-0.76), comparable to the wider level of reproducibility of whole-cell changes (r = 0.09-0.81). The 79 transcripts that changed in abundance after H_2_O_2_ also changed more in nuclear fraction than abundance-matched transcripts that did not, at 15 min in all four cell lines (resampling P ≤ 0.012; **Supplemental Table 5**). However, few individual transcripts passed a false discovery rate of 0.05 (at most 10 per cell line and time point), and the nuclear compartment also contains overlying cytoplasm. We therefore regard these compartment measurements as preliminary. Compartment changes under stress trended in the expected direction, but the nuclear-cytoplasmic partitioning of individual targets cannot yet be measured with confidence.

### An interpretable classifier shows that cell identity, but not exposure time, generalizes across runs

The analyses above tested one target at a time between predefined groups of cells, such as treated and untreated wells or cell-cycle phases. As a complementary approach, we asked how well the complete profile of each individual cell identifies its cell line and treatment, and which features a classifier uses to do so. While the well-level analyses showed that the cell lines differ on average (**Fig. 1D**; **Supplemental Fig. 2**), this approach tests whether single cells can be assigned to the correct cell line. A one-dimensional convolutional neural network trained on all 796 target features assigned untreated cells in a held-out set to the correct cell line with 98.7% accuracy (**Fig. 8A**; **Supplemental Table 6**). Because cells from the same well share well-specific effects, random splits can overstate performance, so we also trained the model on a single run and tested it on every other run. Accuracy remained 93-96% for runs of the same panel version, including runs in a later chemistry period, and 84-89% for the earlier panel version (mean 91.7%; **Fig. 8A**). Multinomial logistic regression and gradient-boosted trees performed similar to the network (held-out-run accuracy 97-99%; **Fig. 8A,B**). Applied to the co-culture wells of the first run, the model trained on run R0418 recovered the same composition as the classifier trained on that run (**Fig. 8C**; **Supplemental Fig. 4D**). Thus, the profile of each single cell encodes its identity robustly and largely linearly, and this identity can be read out across runs, panel versions, and mixed wells.

**Figure 8.**
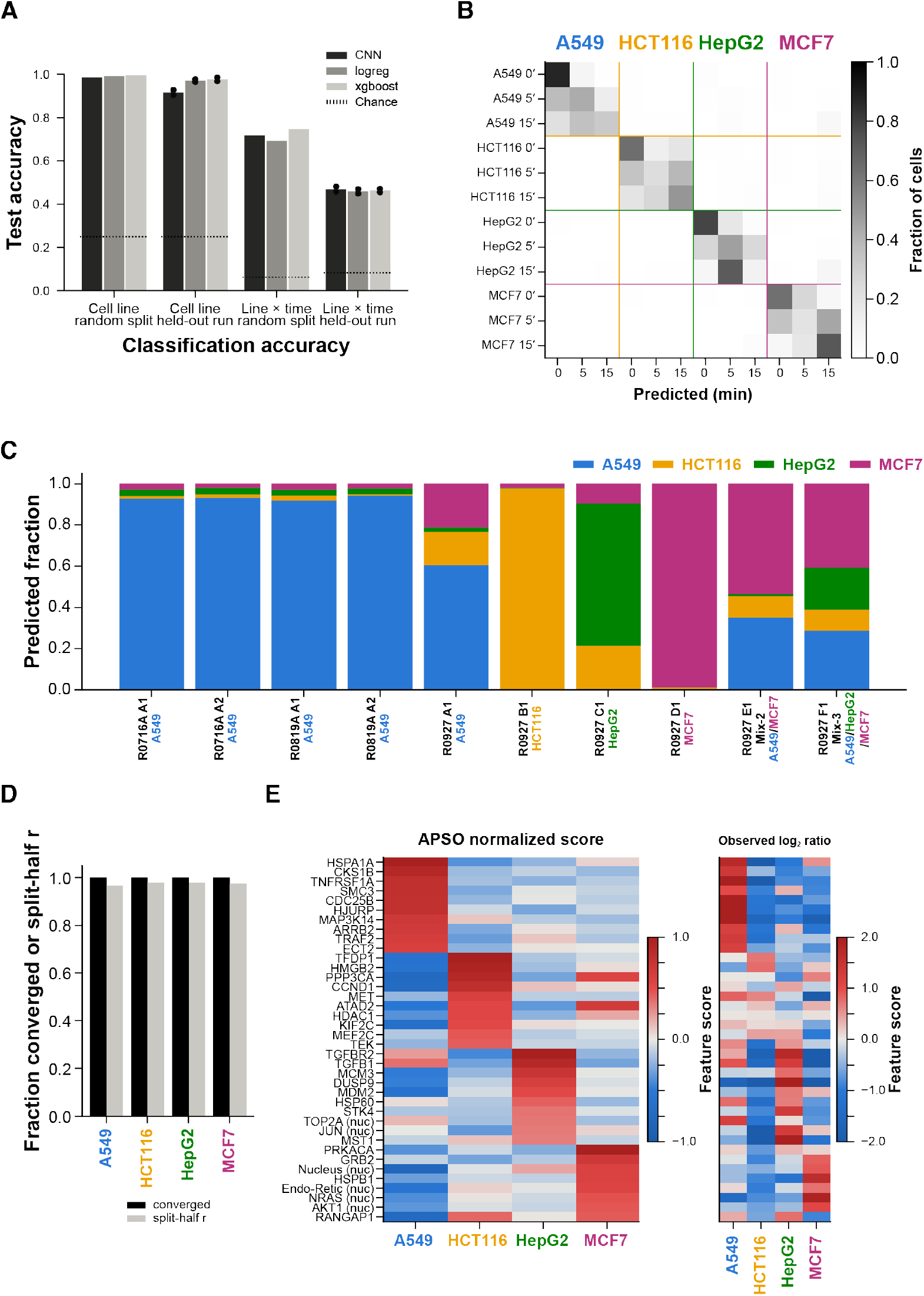
Classifier performance and APSO feature attribution. (A) Test accuracy of the convolutional network (CNN), multinomial logistic regression and gradient-boosted trees (XGBoost) for cell line (untreated cells) and cell line × H_2_O_2_ exposure time, with random splits of cells or with a held-out run (points: the two training runs). Dotted lines indicate chance. (B) Confusion matrix of the held-out-run cell line × time model (both directions pooled; row-normalized). (C) Predicted composition of untreated wells from other runs by the cell-line model trained on run R0418, including the earlier panel version (R0927) and its mixed-cell-line wells, before correction for misclassification. (D) Fraction of swarms converging (global-best probability ≥ 0.9) and split-half correlation of feature importances for each cell line. (E) APSO importance of the ten top-ranked features for each cell line (scaled to a maximum absolute value of 1 within each cell line) alongside the observed log_2_ ratio of each feature between that cell line and the others in real cells.

Exposure time was much harder to predict. Within a run, a 16-class model (cell line × time) reached 71.9% accuracy on a random split. Because each condition occupied a single well per run, however, this accuracy likely reflects differences between wells rather than treatment. When a model for the time points shared by both replicate runs (0, 5 and 15 min; 12 classes) was trained on one run and tested on the other, it still assigned the cell line correctly for 94-98% of cells but assigned exposure time correctly for only 47-52% (chance, 33%; **Fig. 8A,B**). The baseline models performed similarly. Errors were asymmetric: a model trained on R0418 rarely called 15-min A549, HCT116, and HepG2 cells in R0421, consistent with the lower counts per cell of the 15-min wells in R0418 (**Supplemental Fig. 1C**). Thus, individual cells carry little transferable information about exposure time at 5-15 min, and the features that separate treated wells within a run are partly well-and run-specific. Population-level differences remain detectable across thousands of cells, but held-out-run validation, and replicate wells per condition within each run, are needed to separate treatment effects from well effects.

To assess what each measurement type contributes, we repeated the classification with subsets of features (gradient-boosted trees; **Supplemental Table 6**). Cell identity came almost entirely from RNA. Transcripts alone identified the cell line in another run with 98.6% accuracy, close to the 98.9% of all features, and across panel versions with 95.6%. The 48 signaling proteins alone reached 56%, and cell-paint intensities with cell area 77% (chance, 25%). Nuclear-compartment measurements alone retained most of this information (94.5%). Exposure time behaved differently. Within a run, adding protein and morphology features to RNA raised cell line × time accuracy from 68.0% to 72.9%. Across runs, however, no subset of features predicted exposure time well (35-50%; chance, 33%), and adding protein and morphology to RNA did not help (47% vs 50%). We find that RNA carries the cell-identity signal. The within-run gain from protein and morphology probably reflects differences between wells rather than a transferable response to treatment.

We next applied adversarial particle swarm optimization (APSO) to identify the features that the classifier uses to recognize each cell line [44]. APSO generates *in silico* cells that the classifier assigns to a chosen class with high confidence, so the features of these cells reveal what the classifier has learned. For each cell line, we evolved 100 independent swarms of 500 *in silico* cells and attributed each prediction to individual features with integrated gradients (Methods). Every swarm reached a predicted probability of at least 0.9, and feature importances agreed closely between random halves of the swarms (r = 0.97-0.98; **Fig. 8D**). Thus, APSO produced a stable ranking of the features that define each cell line for this classifier.

The highest-ranking features were largely cell-line markers in real data. Of the top-10 features per cell line, 90% were more abundant in that cell line than in the others **(Fig. 8E**; **Supplemental Fig. 6A**), for example ECT2 and *HSPA1A* for A549, *HMGB2* for HCT116, *TGFB1* for HepG2 and *PRKACA* for MCF7 [29, 45]. Five to seven of the ten top-ranked features per cell line were also among the top ten of an independently trained model that used all cells without quality filtering (**Supplemental Table 6**), so APSO attributions are stable across model retraining. APSO applied to the cell line × time model converged for 81-100% of swarms per class (**Supplemental Table 6**). Because exposure-time discrimination did not generalize across runs, however, we do not interpret its time-specific features biologically (**Supplemental Fig. 6B**). The classifiers confirm that cell identity is strongly and reproducibly encoded in the *in situ* profile, mainly by transcripts. In contrast, the effect of a 5-15-min exposure is too small and run-dependent to be learned in a way that transfers between runs.

## Discussion

By measuring transcripts, proteins, phosphoproteins, and cellular morphology in the same cells, we reached three main conclusions about the earliest response to stress, two of which could not have been reached with bulk measurements. Chemically distinct stressors converge on a common early program. They differ mainly in how strongly they engage it, with pathway-specific events such as TNF-α-induced HSP27 phosphorylation added on top. The strength of the response is a property of each cell line rather than of its neighbors, at least within 30 min, because it persisted when weakly and strongly responding cells grew side by side. Cell identity also matters more than cell-cycle state, so the acute response can be studied in asynchronous cells without chemical synchronization, which itself induces stress. These conclusions rest on baseline profiles that were reproducible across runs, panel releases, and chemistry periods and concordant with bulk RNA-seq of the same cell lines.

The response to H_2_O_2_ had two stages, and the first was shared by every stressor we tested. Within the first 15 min, all five stressors changed the same transcripts and phosphoproteins, including increased expression of MAP3K5, which encodes the JNK and p38 activator ASK1 [46, 47], and a modest increase in JNK phosphorylation. The one clear exception, HSP27 phosphorylation after TNF-α, is a known product of TNF-receptor signaling [40, 48]. The assay can therefore detect stressor-specific events when they occur. Because the panel does not include the later effector programs that distinguish these stresses, it may understate their differences. Within the pathways measured, however, the earliest response to stress is largely shared.

The compartment measurements, although preliminary, suggest that the early decreases arise in different ways. For about two-thirds of the transcripts that decreased at 15 min, the decrease was larger in the nucleus than in the cytoplasm, as expected if transcription had slowed. GADD45G instead decreased equally in both compartments, as expected for rapid decay of an unstable transcript. This early loss precedes the induction of GADD45 genes reported after hours of stress [49, 50]. By 30 min, cells had induced the immediate-early genes reported in bulk studies of oxidative stress [19, 37], most strongly in HCT116 and A549. This may reflect differences between the cell lines in antioxidant capacity or MAPK signaling [51]. Thus, H_2_O_2_ first reduces specific signaling transcripts and then induces an immediate-early program whose strength depends on the cell line.

Several of these conclusions required single-cell and spatial measurements. H_2_O_2_ shifted RNA and protein levels in nearly all cells rather than triggering a responsive subset, so population averages describe the typical cell rather than a mixture of responders and non-responders, a distinction that bulk measurements cannot make. Co-culture, which compares cell lines under identical exposure in the same well, showed that the weak immediate-early response of MCF7 cells is intrinsic. It persisted next to strongly responding A549 and HepG2 cells, with no detectable effect of neighbors within 30 min. Because the cell lines share a well, this design also removes the well-to-well variation that limits comparisons between homogenous wells. Even within a homogeneous culture, however, the local environment can matter. Crowded HCT116 cells responded less than cells in sparse regions, possibly because neighboring cells deplete H_2_O_2_ locally [41, 42]. Effective exposure was not measured, but local cell density should be recorded or controlled in single-cell stress experiments.

The cell-cycle analysis also depended on single-cell measurements. Chemical synchronization, the usual method for cell-cycle-resolved bulk measurements, itself activates stress pathways [16, 17]. By inferring cell-cycle position from the same measurements, we could compare the response across the cycle in unperturbed cells. The response was similar in the inferred G1, S and G2/M phases. Within each cell line, the immediate-early fold change varied by at most 1.3-fold across the cycle, far less than between cell lines. A few responses, such as the induction of *JUN* and *DDIT3* in HCT116, were modestly stronger in G2/M cells. The early response also acted on cell-cycle regulators. CDK2 Tyr15 phosphorylation, CHK1 Ser317 phosphorylation and *GADD45G* decreased in every phase. Whether these changes relax checkpoint control, and whether phase-specific consequences such as checkpoint arrest emerge at later times, remain open questions. More generally, the 106 targets that varied along the inferred ordering in four cell lines, together with the DNA-content profile, support the use of targeted *in situ* panels to infer cell-cycle position [24–26, 52].

The classifier analyses also have implications for how *in situ* data should be analyzed. Cell-line identity was learned robustly by the network, by logistic regression, and by gradient-boosted trees, and it transferred across runs and panel releases. APSO attributions for cell identity were also stable across swarms and across independently trained models. Exposure time, in contrast, was predicted well within a run but poorly in a held-out run. Because each condition occupied a single well per run, the within-run accuracy largely reflected differences between wells. Random splits of cells from the same wells can therefore overstate what a classifier has learned about treatment. Held-out-run validation, and replicate wells for each condition within a run, should be standard for *in situ* experiments.

Our analyses also quantify practical aspects of the platform that should inform study design. Ǫuality filters should be evaluated for each cell type: filtering removed a larger fraction of HepG2 cells, which grow in tightly packed colonies (**Supplemental Fig. 1A,B**), so composition estimates were computed from all segmented cells. Signal from abutting cells contributed roughly 1% of a neighbor’s marker level per adjacent cell, an upper estimate that may matter when detecting rare or low-abundance features in heterotypic neighborhoods. In addition, the moderate concordance between transcript and protein abundance for the 18 genes measured both ways indicates that the two modalities provide partly independent information.

These results suggest several directions for future work. Extending the time course beyond 30 min and across a range of H_2_O_2_ doses will test how the shared early program resolves and how the immediate-early response is followed by stressor-specific effector programs. Panels that include antioxidant, metal-response, unfolded-protein and inflammatory genes would capture these programs. Co-culture of cell types that interact *in vivo*, such as epithelial cells with immune or stromal cells, will test whether the cell-intrinsic behavior we observed between cancer cell lines extends to cell types that communicate through paracrine signals. Applying the same approach to primary cells and tissue sections will also test how these observations apply in more physiological contexts.

The earliest response of the MAPK and cell-cycle regulators measured here is largely shared among chemically distinct stressors in A549 cells and differs in magnitude among cell lines. These cell-line differences persist when cell lines share the same well and are larger than the differences among inferred cell-cycle phases. Single-cell *in situ* measurements made it possible to test these properties in co-culture and without synchronization, and the analysis framework we provide can be applied to other perturbations, cell types and tissues.

## Methods

### Cell culture

A549 (ATCC CCL-185), HCT116 (ATCC CCL-247), HepG2 (ATCC HB-8065) and MCF7 (ATCC HTB-22) cells were cultured in Iscove’s Modified Dulbecco’s Medium with L-glutamine and 25 mM HEPES (Corning 10-016-CM) supplemented with 10% fetal bovine serum, 2 mM L-glutamine and 1% penicillin-streptomycin, at 37 °C in 5% CO_2_. Cells were used within 20 passages and tested negative for mycoplasma. For cytoprofiling, cells were seeded directly into the wells of AVITI24 flow cells at 10,000-20,000 cells per well and grown for 20-36 h before treatment. Co-culture wells were seeded with A549 and MCF7 (Mix-2) or A549, MCF7 and HepG2 (Mix-3) at ratios intended to yield equal proportions at the time of the experiment.

### Oxidative stress and additional stressor treatments

H_2_O_2_ (30% (w/w) stock; Sigma-Aldrich H1009) was freshly diluted in pre-warmed complete medium and added to a final concentration of 300 µM by replacing the culture medium for 5, 15 or 30 min at 37 °C. Untreated control wells were not exposed to H_2_O_2_ but underwent a mock replacement with fresh medium. Exposure was terminated by aspirating the medium, two brief PBS washes and immediate fixation. In the multi-stressor runs, A549 cells were exposed for 5 or 15 min to H_2_O_2_ (300 µM), recombinant human TNF-α (100 ng/mL), cadmium chloride (200 µM), dithiothreitol (10 mM) or lipopolysaccharide (*E. coli* O111:B4, 10 µg/mL) premixed in fresh media. Mock-treated wells also underwent a mock media replacement without a stressor.

### AVITI24 *in situ* cytoprofiling

Cells were processed with the AVITI24 Teton CytoProfiling workflow using the Teton MAPK/Cell Cycle panel (Element Biosciences) according to the manufacturer’s instructions. Run R0927 used a pre-production version of the commercial panel whose chemistry and reagents were identical except for two antibodies. Briefly, cells were fixed with 4% paraformaldehyde for 15 min at room temperature and permeabilized, stained with cell paint reagents labeling the plasma membrane, nucleus, F-actin, endoplasmic reticulum, Golgi apparatus and mitochondria (wheat germ agglutinin, a far-red DNA stain, phalloidin, calreticulin, GM130 and a mitochondrial 16S rRNA probe in the first panel release), and incubated with target-specific RNA probes and DNA-barcoded antibodies against 48 protein and phosphoprotein epitopes. Runs were performed on three AVITI24 instruments (AV224504, AV224101 and AV234001) between September 2024 and August 2025 (**Supplemental Table 1**).

### Image processing and per-cell feature extraction

Images were processed with the onboard AVITI24 analysis software, which segments nuclei and cells, assigns decoded RNA and protein barcodes to the whole-cell and nuclear compartments, quantifies cell paint intensity and morphology features, and reports per-cell tables (RawCellStats) that were used for all analyses. Each well was imaged as 12 tiles, and cell coordinates are reported within tiles (0.48 µm per pixel in all runs, as derived from the paired pixel and micrometer coordinates in the data files of the panel-v2 runs).

### Panel harmonization and feature construction

Per-cell RawCellStats tables exported from each run were used for all downstream analyses. The first run (0 and 30 min) was performed with an earlier release of the Teton MAPK/Cell Cycle panel that differed from later runs in barcoding-batch assignment and in antibody and stain nomenclature. Features from this run were harmonized to the later panel by target name: RNA batch indices were offset by one, renamed phospho-epitope antibodies were mapped to their updated names, cell paint channels were mapped to their compartment names (WGA, cell membrane; RedDot, nucleus; phalloidin, actin; calreticulin, endoplasmic reticulum; GM130, Golgi apparatus; 16S, mitochondria), and antibodies present in only one panel release were excluded (**Supplemental Table 1**). Unassigned and non-specific-binding decoy channels were not used. The resulting feature set comprised 796 features per cell: whole-cell and nuclear measurements for 48 protein epitopes (23 phospho-specific), 344 RNA probe sets (338 transcripts and a GAPDH control in each of six imaging batches) and six cell paint stains. We refer to each transcript, protein or stain as a target, and to its measurement in one compartment as a feature. Every RNA and protein feature had nonzero counts in every well of the time-course runs, with at least 259 counts per well for each whole-cell transcript, so no feature was excluded for low counts. Analyses that required a minimum count state the threshold.

### Classifier input normalization

For the original and random-split classification, cells from the three H_2_O_2_ time-course runs were pooled (mixed-cell-line wells excluded) and each feature was divided by its maximum value across the cells used for that model, placing every feature on a [0, 1] scale. Missing values were set to zero. Cells were labeled by cell line and exposure time (16 classes) or, for the cell-line model, restricted to untreated cells (4 classes).

### Cell quality control

Cells were retained when (i) a nucleus had been segmented (non-zero nuclear DNA-stain signal), (ii) log10 cell area and (iii) log10(total RNA counts + 1) fell within three scaled median absolute deviations of the median of their well, and (iv) total RNA counts were at least 10. Thresholds were computed per well because the cell lines differ in size and RNA content. Per-well cell numbers before and after filtering are reported in **Supplemental Table 1** and **Supplemental Fig. 1**. Unless otherwise stated, analyses use ǪC-passing cells. The classifier and APSO analyses were additionally run on all cells without quality filtering.

### Pseudobulk differential abundance

For each well, per-cell RNA and protein counts (whole-cell compartment) were summed across ǪC-passing cells. For this and all other per-target analyses, the six GAPDH control probe sets (one per imaging batch) were summed into a single GAPDH target. Differential abundance was tested separately for each cell line and modality with PyDESeq2 (v0.4.12) using the design ∼ run + time, so that every exposure time was compared with untreated cells from the same run while run-level differences in chemistry, imaging and cell density were absorbed by the run term. Size factors were estimated per modality by the median-of-ratios method, and P-values were adjusted by the Benjamini-Hochberg procedure. Targets with adjusted P < 0.05 were considered significant. For A549, an extended model additionally included the H_2_O_2_ and mock wells of the two multi-stressor runs, giving four independent runs at 5 and 15 min. Targets significant in at least one cell line × time contrast were clustered by Ward linkage on their log2 fold-change profiles. Median-of-ratios size factors assume that most targets do not change between conditions. This held here since for any single comparison of the time-course model, at most 11% of transcripts and 27% of proteins changed significantly, and in the extended A549 model at most 32% of transcripts changed. No single control gene, such as GAPDH, was used for normalization. Per-cell scores, such as the IEG score, were expressed per 10⁴ RNA counts of the cell. This removes differences in total detected RNA between cells, with the caveat that it would also remove a genuine global change in transcription.

### Reproducibility and RNA-protein concordance

Replicate-run concordance was computed as the squared Pearson correlation of log10 per-well mean counts per cell (ǪC-passing cells) between matched wells of the two replicate time-course runs, separately for RNA and protein targets. For well-to-well comparisons, summed RNA counts per well were converted to log2 counts per million and correlated (Pearson), followed by average-linkage hierarchical clustering on correlation distance. Identity of wells from the first (panel-v1) run was assessed by correlating their run-centered log2 CPM profiles with run-centered cell-line centroids from the two replicate runs. For RNA-protein concordance, the 18 targets measured by both a probe set and an antibody were normalized within modality (CPM), averaged across the two replicate runs, log2-transformed and z-scored across the four cell lines.

### Nuclear fraction and compartment-resolved changes

For each well and target, the nuclear fraction was the sum of nuclear-compartment counts divided by the sum of whole-cell counts over ǪC-passing cells. Nuclear enrichment was the nuclear fraction divided by the median ratio of nuclear to cell area in that well (panel-v2 runs). Expected localization of proteins was assigned from UniProt and the Human Protein Atlas [53, 54]. Changes in nuclear fraction were modeled per cell line and target as logit(nuclear fraction) ∼ run + time by least squares on wells (targets with ≥20 counts in every well). To separate target-specific redistribution from well-wide shifts in the nuclear share of counts, the logit nuclear fractions of each well were first centered on their median across targets of the same modality, and uncentered estimates are reported alongside. Residual variances were moderated across targets by an empirical-Bayes prior fitted by the method of moments, as in limma, and P-values were BH-adjusted within each cell line and contrast [55]. Compartment-resolved fold changes in A549 cells were computed per run from nuclear and cytoplasmic (whole minus nuclear) counts, each divided by the well total RNA count, and averaged over runs. To benchmark the nuclear compartment, the nuclear share of each cell-paint stain was divided by the nuclear share of cell area in every ǪC-passing cell of the panel-v2 runs, and medians were taken per run. Replicate-run reproducibility was the Pearson correlation, across RNA targets, of the well-median-centered logit nuclear fraction of matched wells of the two replicate runs, and of its change from the untreated well of the same run. Responsive transcripts (FDR < 0.05 in any cell line and time point) were compared with 2,000 sets of non-responsive transcripts drawn with replacement within deciles of mean abundance, using the median absolute change in centered logit nuclear fraction.

### Cell-cycle inference

For each cell line, per-cell RNA counts (GAPDH probes excluded) were normalized to 10,000 counts and log1p-transformed. S and G2/M scores were computed within each run as the mean expression of the S-phase (40 genes) and G2/M (53 genes) sets of Tirosh et al. (2016) minus the mean of expression-matched control genes (25 expression bins), with controls drawn only from non-cell-cycle targets because the panel is enriched for cell-cycle genes [25]. Phases were assigned with the scanpy rule (G1 if both scores < 0, otherwise the higher-scoring phase) [56]. The 93 cell-cycle genes were z-scored within each run, a two-component PCA was fitted on untreated cells and all cells were projected. The angle of each cell around the centroid was rotated so that the circular mean of G1 cells was zero and oriented so that S preceded G2/M, then converted to a cell-cycle position in [0, 1) by the empirical cumulative distribution of untreated cells. DNA content was the integrated DNA-stain signal within the nuclear mask divided by the median of G1 cells in the same well. Periodic targets were identified from the fraction of variance explained by 20 position bins (median η² ≥ 0.03 across cell lines, untreated cells, run-centered log-normalized expression), and their z-scored bin profiles concatenated across cell lines were clustered by k-means (k = 5).

### Cell-cycle-resolved response

Cells were assigned to G1 (position ≥ 0.85 or < 0.15), S (0.15–0.5) or G2/M (0.5–0.85) phases, chosen from the score and DNA-content profiles. Counts were summed per well and phase, and differential abundance was tested within each phase as above. Phase dependence of each target’s response was tested with Cochran’s Ǫ statistic on the three phase-specific log_2_ fold changes and their standard errors (χ² with 2 degrees of freedom), BH-adjusted within each cell line and contrast. This test ignores covariance between phases and can be conservative if their errors are positively correlated. For a more sensitive within-well comparison, log_2_ CPM of each target was computed per well and phase, and the difference between two phases of the same well was modeled as ∼ run + time with empirical-Bayes moderated variances, so that the time coefficient estimates the difference between the phase-specific log_2_ fold changes. For variance decomposition (**Supplemental Table 4**), the cell line × phase table of log_2_ fold changes of each responsive target was partitioned by two-way analysis of variance into cell-line, phase, and residual sums of squares.

### Neural-network classifier

A one-dimensional convolutional network (three convolutional layers with 16, 32 and 64 channels, kernel sizes 5, 5 and 3, ReLU activation and average pooling, followed by a 128-unit fully connected layer with dropout 0.3) was trained on per-cell feature vectors scaled by their feature-specific maxima with class-balanced cross-entropy, Adam (learning rate 10⁻⁴) and mini-batches of 64 cells, selecting the epoch with the highest validation accuracy (maximum 250 epochs for cell line × time and 100 for cell line; early stopping after 30 epochs without improvement). Cells were split into training (80%), validation (10%) and test (10%) sets, stratified by class. For held-out-run evaluation, models were trained and validated on one replicate run and tested on the other (and, for the cell-line model, on all other runs), with per-feature scaling computed from the training run only. Multinomial logistic regression (L2, class-balanced) and gradient-boosted trees (XGBoost, histogram method, up to 400 rounds, depth 6, early stopping on the validation set) were fitted to the same splits as baselines, on stratified subsamples of up to 200,000 (logistic regression) or 300,000 (XGBoost) training cells.

### Adversarial particle swarm optimization and attribution

Class-specific *in silico* cells were generated by adversarial particle swarm optimization against the trained classifier: 500 particles were initialized as uniform random vectors with 95% of entries set to zero, and updated for 20 iterations with inertia 0.8 and cognitive and social coefficients 0.5 (Clerc–Kennedy constriction applied when their sum exceeds 4), with each particle rescaled to [0, 1] after every step, to maximize the softmax probability of the target class [44]. After the final step, integrated gradients (zero baseline, 50 steps; Captum) were computed for every particle [57]. This was repeated 100 times per class with different random seeds. For each run, particle positions and integrated-gradient attributions were averaged over particles, and feature importance for a class was the mean over runs of the product of the averaged position and the averaged attribution. Runs were vectorized on a GPU, and equivalence with the original per-particle implementation was verified step-for-step with identical random seeds.

### Comparison with bulk RNA-seq

Normalized expression (nTPM) of the four cell lines and of all 1,206 cell lines of the Human Protein Atlas cell-line atlas (rna_celline.tsv) was log_2_(nTPM + 1)-transformed. AVITI24 RNA counts of untreated wells of panel-v2 runs (ǪC-passing cells) were summed per well, converted to CPM, averaged per cell line and log_2_(CPM + 1)-transformed, excluding GAPDH probes. Absolute concordance was the Spearman correlation across panel genes. Relative concordance used per-gene z-scores across the four cell lines. For identity matching, the AVITI24 per-gene z-scores of each line were correlated (Pearson) with the per-gene z-scores of every HPA cell line computed across all 1,206 lines.

### Modality contributions

Gradient-boosted tree classifiers (XGBoost; 200 rounds, depth 6) were trained on ǪC-passing cells using subsets of features (whole-cell RNA, whole-cell protein, cell paint (six stains in whole-cell and nuclear compartments) with cell area, RNA + protein, nuclear compartment only, whole-cell compartment only, or all features), scaled by their maxima in the training data, for cell line (untreated cells; train R0418, test R0421 or R0927), cell line × time (random 80/20 split of R0418 and R0421) and exposure time within each cell line (train R0418, test R0421).

### Co-cultured wells

Cells of the mixed wells of run R0927 were assigned to cell lines with a gradient-boosted tree classifier trained on the pure wells of the same run (untreated and 30-min wells; 80/20 split for evaluation). For the IEG score analyses (**Fig. 4B,D**), a second classifier of the same design excluded the five immediate-early probes in both compartments. For the comparison of all targets between mixed and pure wells (**Fig. 4C**), a third classifier of the same design used all features. Summed counts of its high-confidence cells (classifier probability > 0.95) were converted to CPM within modality for each cell line and context, log_2_ fold changes (30 vs 0 min) were computed with a pseudocount of 10 CPM, and standard errors were obtained by bootstrapping cells within wells (200 replicates). Because each context is a single well, a target was called co-culture-dependent only if it differed from the pure well in the same direction in both mixed contexts by more than the 99th percentile of the differences between the two mixed contexts, with bootstrap |z| > 4 in each. To estimate signal spillover without circularity, 15 A549-enriched and 15 MCF7-enriched probes (every other probe of the 30 most differential in each direction, both compartments) were excluded from the classifier and summed (per 10⁴ RNA counts) as held-out marker scores. Observed predicted fractions were corrected for misclassification by non-negative least squares using the classifier’s confusion matrix on held-out pure-well cells. Composition was computed on all cells because ǪC removes most HepG2 cells of this run. Neighbors were identified within each imaging tile (cell coordinates are tile-local). Homotypic clustering compared the fraction of same-type cells among the 10 nearest neighbors with that after random relabeling within the tile. Spillover was the slope of the held-out marker score of high-confidence cells (classifier probability > 0.95) in Mix-2 wells on the number of heterotypic cells among the six nearest neighbors, divided by the mean marker score of the neighboring cell type. Immediate-early fold changes (**Supplemental Fig. 4B**) were computed from total-count-normalized mean expression of cells with classifier probability > 0.9. The immediate-early gene (IEG) score of each cell was the sum of FOS, JUN, DDIT3, DUSP1 and NR4A1 counts per 10⁴ RNA counts. To bound stress-induced transfer between neighbors, the IEG score of high-confidence MCF7 cells was modeled as well + n + n × treated, where n is the number of A549 cells among the six nearest neighbors (capped at five), with standard errors clustered by imaging tile. The upper 95% confidence bound of the interaction was expressed relative to the difference between the mean A549 and MCF7 responses in the same wells. Robustness was assessed with 3 or 10 nearest neighbors, uncapped counts, the A549 fraction among neighbors, counts within 20, 30 or 50 µm, Gaussian-weighted exposure (σ = 15 or 25 µm), a mixed model with a random intercept per tile, and a null distribution from 1,000 permutations of cell-line labels within tiles.

### Local cell density

For ǪC-passing cells of the pure wells of the time-course runs, local density was computed from all segmented cells of the same imaging tile as the negative logarithm of the distance to the tenth nearest neighbor, excluding cells within 50 µm of a tile border, and converted to quintiles within each well. The early-response score was the sum of counts of seven transcripts that decreased at 15 min in at least three cell lines (GADD45G, FLT4, MAPK8IP1, MAPK8IP2, MAPK8IP3, SOS1 and NCAPD2) per 10⁴ RNA counts. Fold changes versus untreated cells of the same run were computed from counts pooled within each quintile, with 95% confidence intervals from resampling tiles within wells. Adjusted effects were estimated from the cells of the most and least crowded quintiles, per cell with tile fixed effects and covariates for log cell area, cell-cycle position (sine and cosine) and log total RNA, with standard errors clustered by tile. In this model, the early-response score is in standard-deviation units and the IEG score was log-transformed (log_2_ units).

### Software and statistics

Analyses were performed in Python 3.11 with pandas 2.2.3, NumPy 1.26.4, SciPy 1.14.1, statsmodels 0.14.4, PyDESeq2 0.4.12, scikit-learn 1.5.2, XGBoost 2.1.2, PyTorch 2.5.1, Captum 0.7.0 and matplotlib 3.9.2. Statistical tests were two-sided unless stated otherwise, and P-values were adjusted by the Benjamini–Hochberg procedure within each family of tests as defined in the corresponding section. Numbered shell scripts that reproduce every analysis, figure panel and supplemental table, together with the software environment, are provided as Supplemental Code.

## Data access

Per-cell RawCellStats tables and curated metadata will be deposited at Zenodo upon publication. Analysis code that reproduces all figures and tables is provided as Supplemental Code and at https://github.com/EpiGenomicsCode/2026_AVITI24.

## Acknowledgments

Sample processing and FAIR-compliant analysis were made possible by the Platform for Epigenomic and Genomic Research (RRID:SCR_021861) and Cornell University Biotechnology Resource Center Epigenomics Core Facility (RRID:SCR_021287). This work was supported by the National Institutes of Health (National Institute of General Medical Sciences) grants R35GM145217, R01ES034353 to BFP, and R35GM155380 to WKML. Computational resources were provided by NSF-ACCESS (BIO220026) to WKML.

## Author contributions

WKML conceived the study. AMG, CMS, and AML performed cell culture, treatments, and AVITI24 cytoprofiling. OWL and WKML analyzed the data and developed the analysis code. BFP and WKML supervised the work and acquired funding. OWL and WKML wrote the manuscript with input from all authors.

## Supplemental figure legends

**Supplemental Figure 1.**
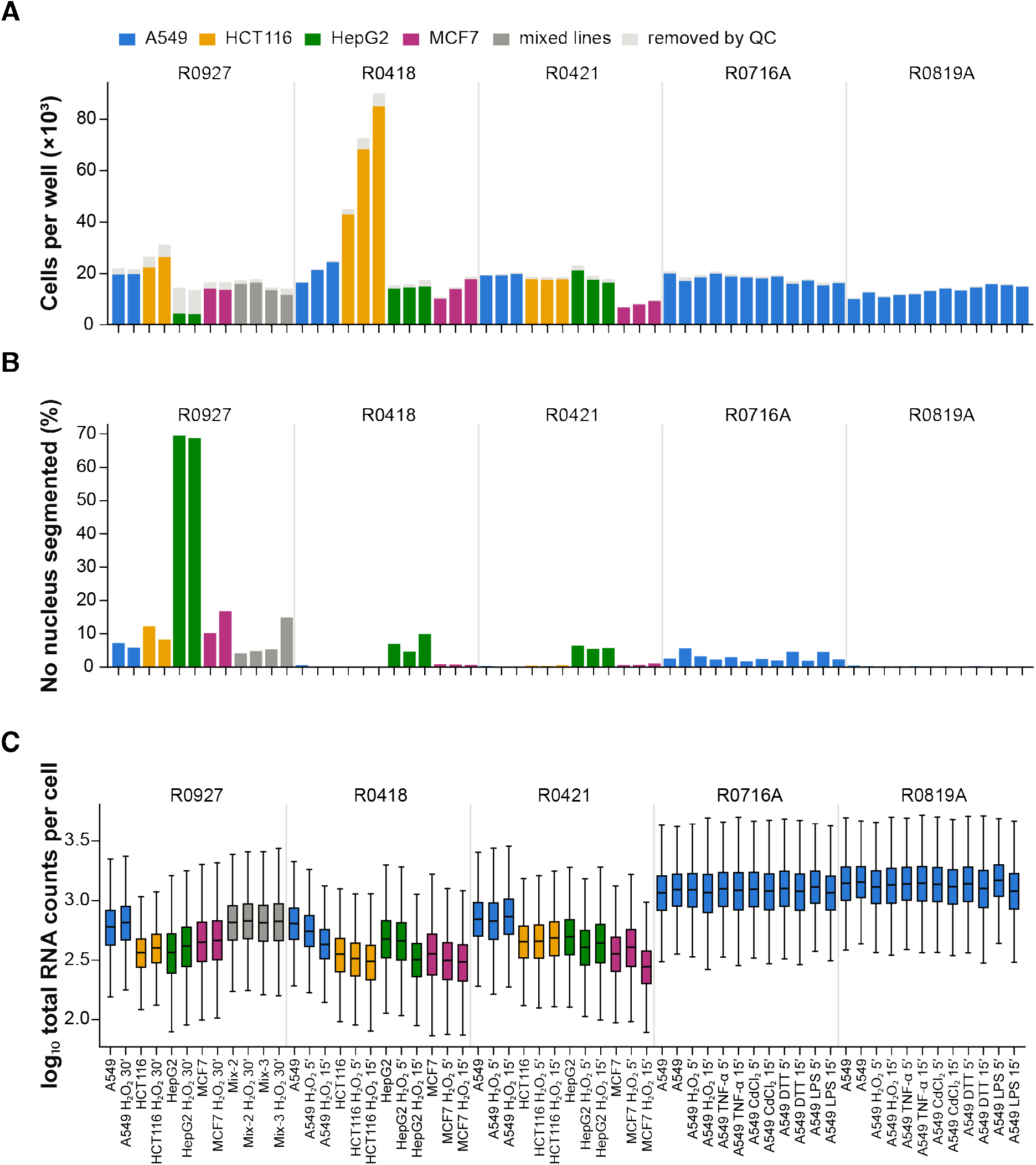
Ǫuality control. (A) Cells per well before (light gray, removed by ǪC) and after ǪC. (B) Percentage of cells without a segmented nucleus. (C) Total RNA counts per ǪC-passing cell.

**Supplemental Figure 2.**
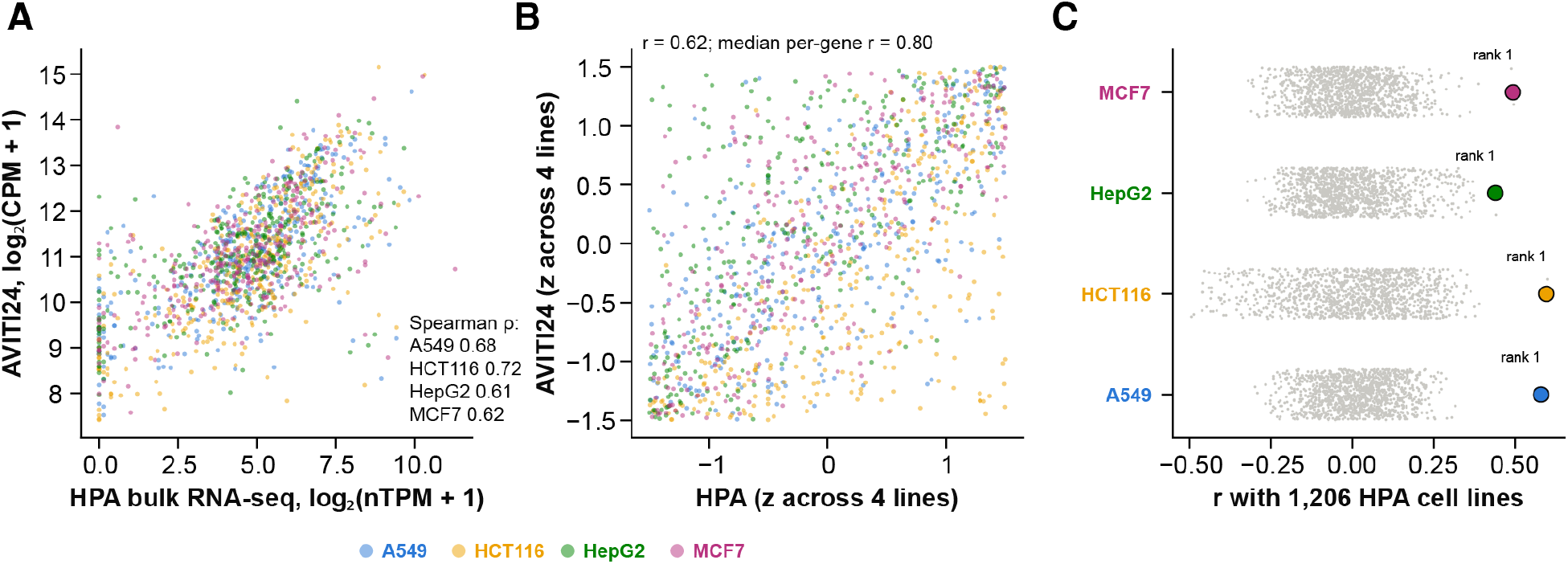
Orthogonal validation against bulk RNA-seq (Human Protein Atlas cell-line atlas). (A) AVITI24 log_2_ CPM versus bulk log_2_(nTPM + 1) for panel transcripts in each cell line. (B) Per-gene z-scores across the four cell lines. (C) Correlation of each AVITI24 cell-line profile with all 1,206 cell lines in the atlas (gray) and with the matching line (colored).

**Supplemental Figure 3.**
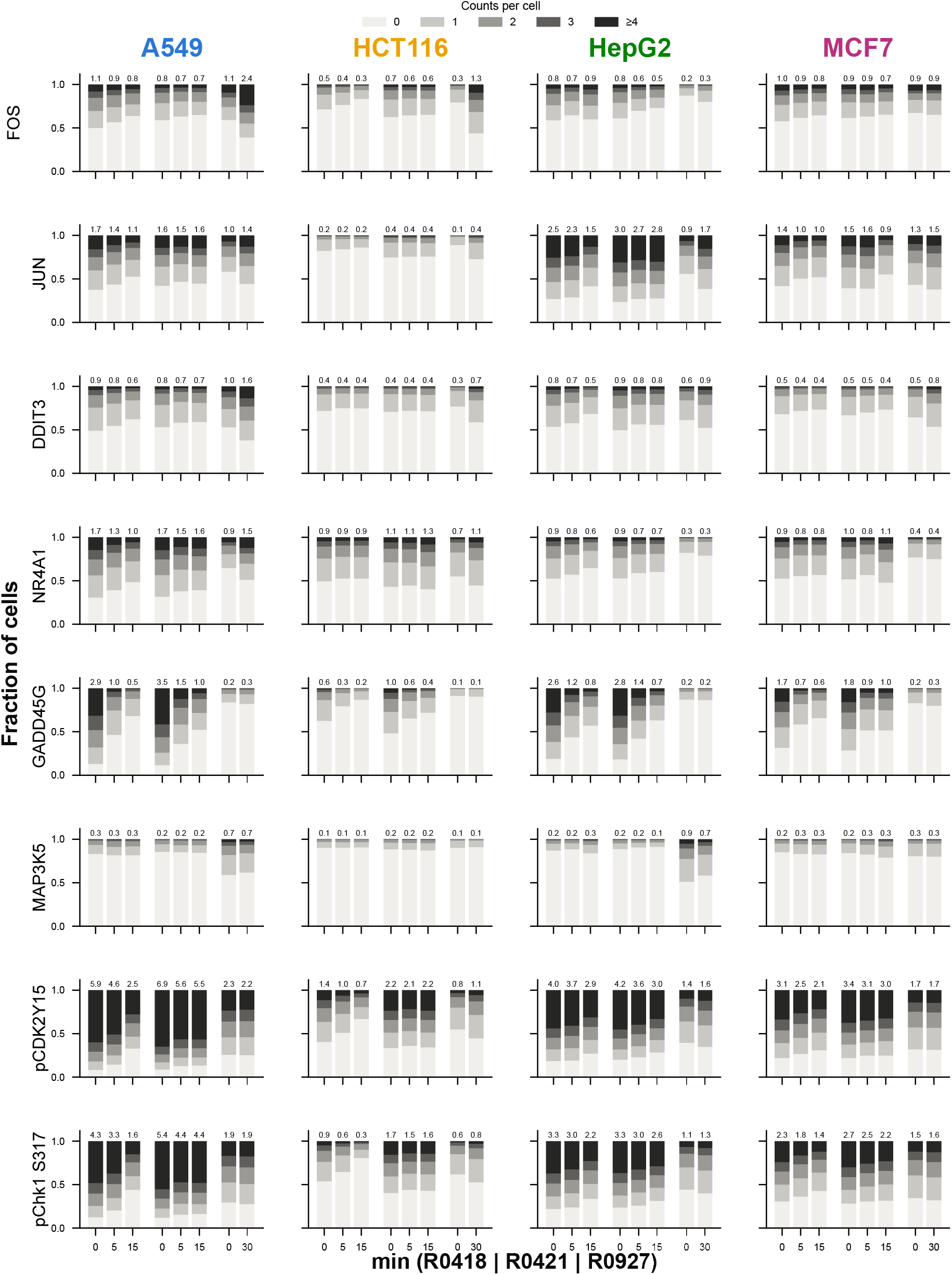
Per-cell distributions of exemplar targets. For each target and cell line, the fraction of ǪC-passing cells with 0, 1, 2, 3 or ≥4 whole-cell counts in each well, grouped by run (R0418 and R0421: 0, 5, 15 min; R0927: 0, 30 min) so that each exposure is compared with untreated cells of the same run. Numbers above bars are mean counts per cell.

**Supplemental Figure 4.**
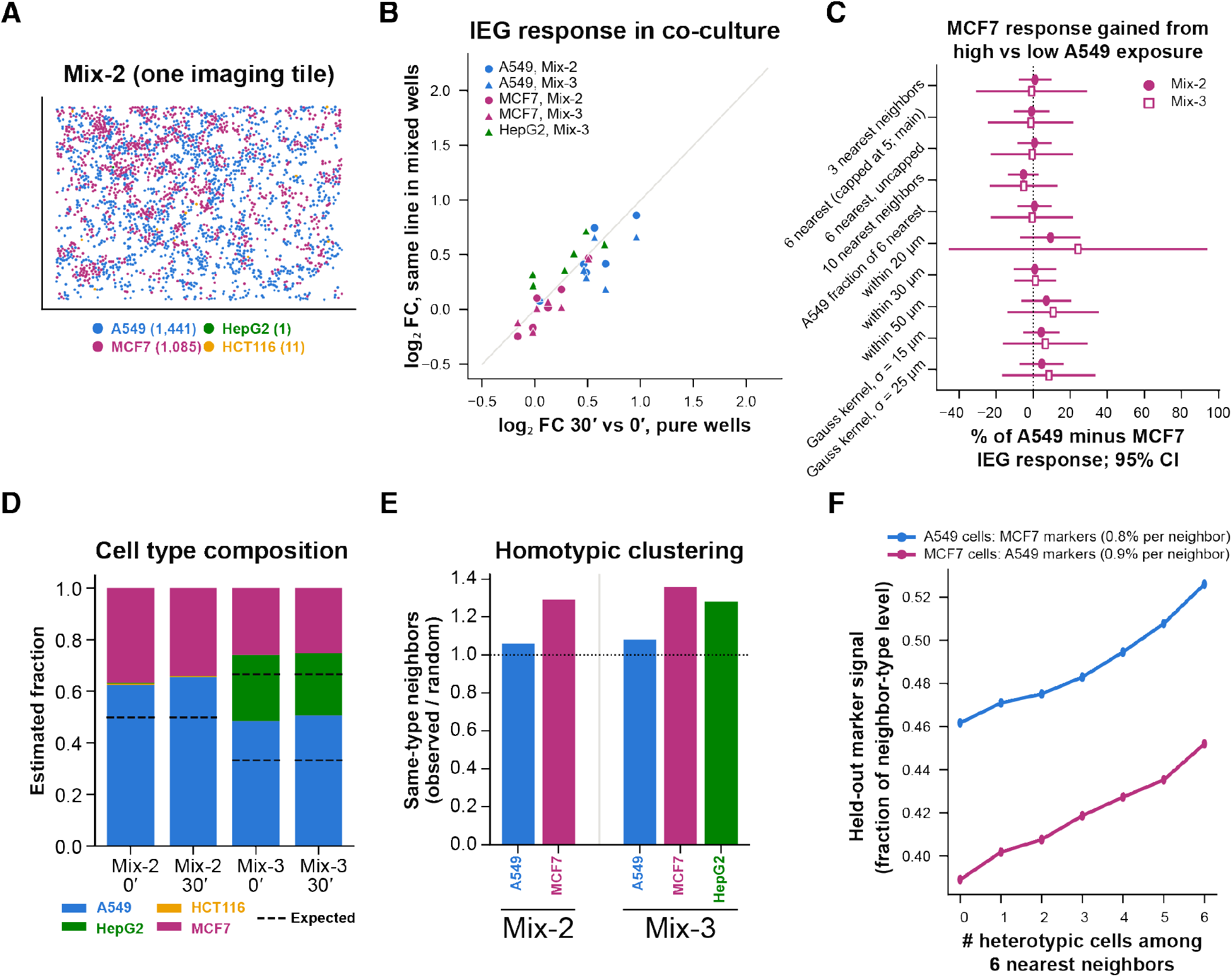
Co-culture quality metrics. (A) One imaging tile of a Mix-2 well colored by predicted cell line. (B) Immediate-early fold changes in mixed versus pure wells. (C) Upper bound on stress-induced transfer of the immediate-early response to MCF7 cells: gain in MCF7 response with high versus low A549 exposure, as a percentage of the difference between the A549 and MCF7 responses, for ten neighborhood definitions (95% confidence intervals; Mix-2 and Mix-3). (D) Composition of the Mix-2 and Mix-3 wells estimated from all cells and corrected for classifier misclassification; dashed lines show the intended proportions. (E) Same-type neighbors relative to random relabeling within tiles. (F) Signal of held-out marker probes of the other cell line versus the number of heterotypic neighbors (signal spillover).

**Supplemental Figure 5.**
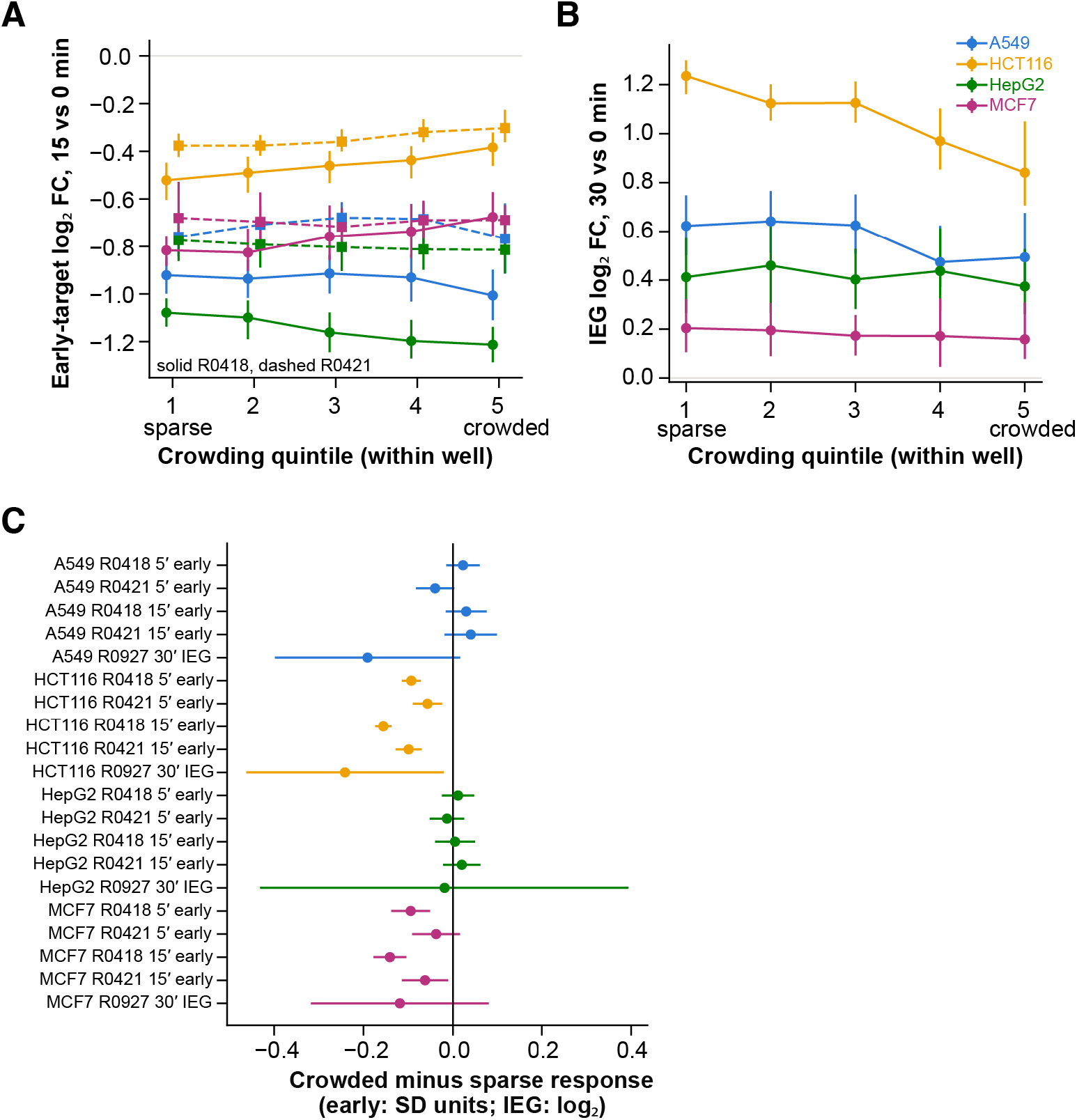
Response to H_2_O_2_ by local cell density in pure wells. (A) Log2 fold change of the early-decrease score (seven transcripts; 15 vs 0 min) by quintile of local density within each well, for each cell line and time-course run. (B) Log2 fold change of the immediate-early gene score (30 vs 0 min) by density quintile. (C) Difference in response between the most and least crowded quintiles, adjusted for cell area, total RNA and cell-cycle position, for every cell line, run and contrast (early-decrease score in standard-deviation units, immediate-early score in log2 units; 95% confidence intervals).

**Supplemental Figure 6.**
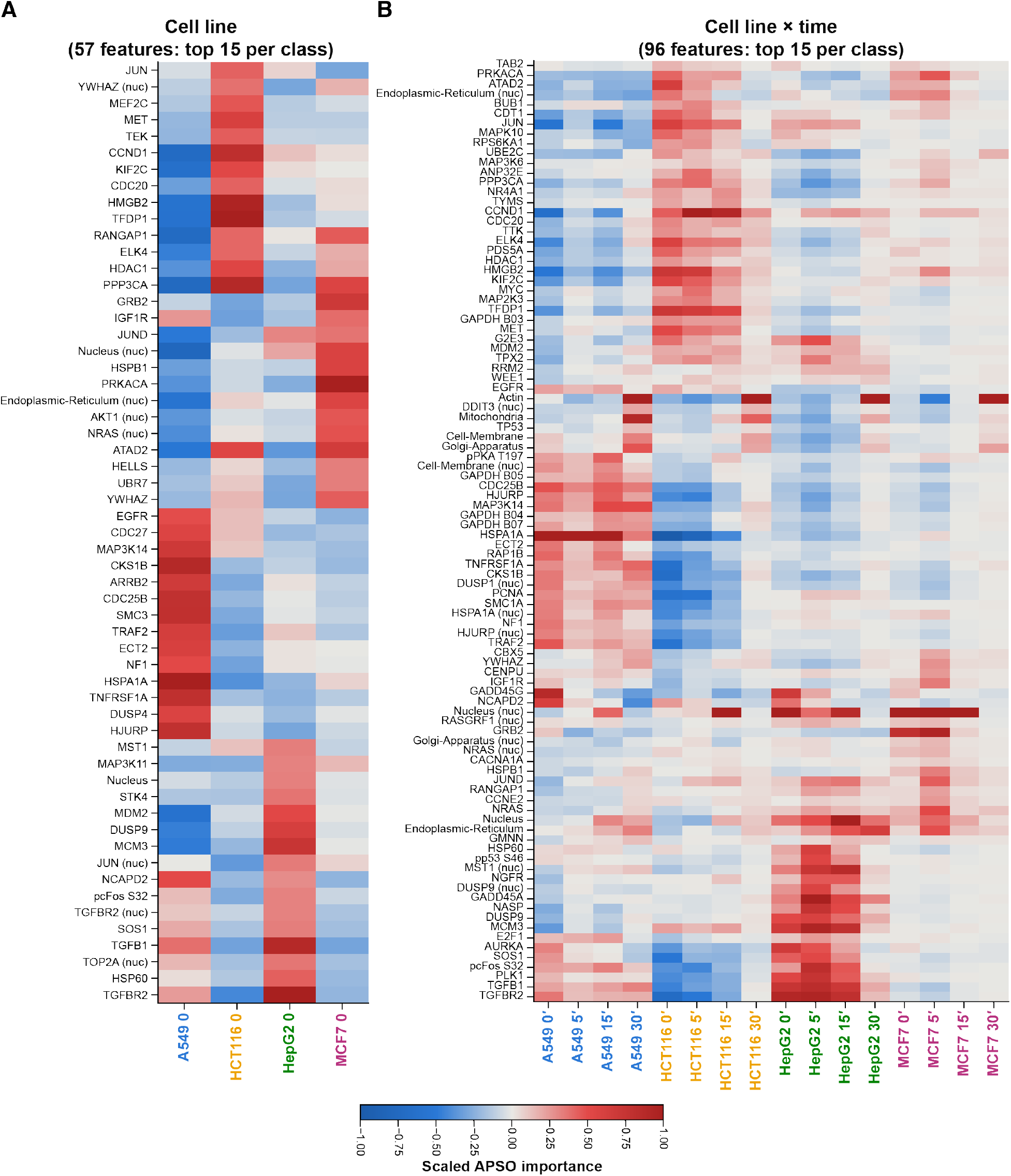
APSO feature importance for every class of the cell-line (A) and cell line × time (B) models (ǪC-passing cells): union of the 15 highest-ranked features per class, scaled within each class and hierarchically clustered. In the cell line × time model, the 30-min classes, which derive from a single run with an earlier panel version, are dominated by cell paint intensity features, consistent with run-and panel-specific rather than treatment-specific discrimination.

## Supplemental table legends

**Supplemental Table 1.** Experimental design, data quality and validation: curated per-well annotation of all runs, harmonization of panel release 1 to release 2 feature names, the 796 per-cell features, per-well detection of every feature, per-well cell numbers before and after ǪC, replicate-run R² per cell line, time point and modality, well-to-well RNA profile correlations, identity of panel-v1 wells, RNA and protein z-scores for paired targets, and comparison with bulk RNA-seq of the Human Protein Atlas (absolute and relative concordance, per-gene correlations and identity matching against 1,206 cell lines).

**Supplemental Table 2.** Response to H_2_O_2_ and to four additional stressors: pseudobulk differential abundance for every target, cell line and time point (ǪC-passing and all cells; A549 extended model), the number and fraction of targets significant in each comparison, target clusters, and per-cell count distributions and Fano factors of exemplar targets with the values expected for uniform and subpopulation responses, for the H_2_O_2_ time course; differential abundance versus mock for A549 cells exposed to five stressors, pairwise correlation between conditions, loadings on the shared response axis, the magnitude of each well along it, and phospho-JNK relative to total JNK.

**Supplemental Table 3.** Co-cultured wells and local cell density: accuracy of the cell-line classifiers, immediate-early response by cell line and context, neighbor analysis and bounds on stress-induced transfer between neighbors, mixed-versus-pure comparisons, composition, homotypic clustering, signal spillover, and response by local cell density in pure wells.

**Supplemental Table 4.** Cell-cycle inference and response by cell-cycle state: PCA loadings of S and G2/M genes, association with inferred cell-cycle position for every target, comparison of the 93 reference genes with the periodic targets and with the H2O2 response, phase-resolved differential abundance, heterogeneity tests, phase occupancy, immediate-early fold change across ten cell-cycle bins, within-well comparisons of phases, equivalence tests and variance decomposition.

**Supplemental Table 5.** Subcellular partitioning: nuclear enrichment in untreated cells, changes in nuclear fraction after H_2_O_2_, compartment-resolved fold changes in A549 cells, nuclear-compartment enrichment of the cell-paint stains, replicate-run reproducibility of nuclear-fraction changes, and comparison of responsive with abundance-matched transcripts.

**Supplemental Table 6.** Classifier performance, modality contributions and APSO feature attribution: test accuracy and macro-F1 of the CNN, logistic regression and gradient-boosted trees for every split, held-out-run exposure-time classification for each training run, predicted composition of wells from other runs, accuracy of classifiers trained on subsets of features, APSO top features, swarm convergence and stability and feature importance for the cell-line and cell line × time models (ǪC-passing and all cells).

